# Structural basis and pathological implications of the dimeric OS9-SEL1L-HRD1 ERAD Core Complex

**DOI:** 10.1101/2025.06.13.659592

**Authors:** Liangguang Leo Lin, Emir Maldosevic, Linyao Elina Zhou, Ahmad Jomaa, Ling Qi

**Affiliations:** Department of Molecular Physiology and Biological Physics, University of Virginia, School of Medicine, Charlottesville, VA 22903, USA; Department of Biochemistry and Molecular Genetics, University of Virginia, School of Medicine, Charlottesville, VA 22903, USA

**Keywords:** ER-associated degradation, core complex, Cryo-EM structure, dimers, disease variants, crosslinking, mutagenesis, protein-conducting channels

## Abstract

The SEL1L-HRD1 complex represents the most conserved branch of endoplasmic reticulum (ER)-associated degradation (ERAD), a critical pathway that clears misfolded proteins to maintain ER proteostasis. However, the molecular organization and pathogenic mechanisms of mammalian ERAD have remained elusive. Here, we report the first cryo-EM structure of the core mammalian ERAD complex, comprising the ER lectin OS9, SEL1L, and the E3 ubiquitin ligase HRD1. The structure, validated by mutagenesis and crosslinking assays, reveals a dimeric assembly of the core complex in which SEL1L and OS9 form a claw-like configuration in the ER lumen that mediates substrate engagement, while HRD1 dimerizes within the membrane to facilitate substrate translocation. Pathogenic SEL1L mutations at the SEL1L-OS9 (Gly585Asp) and SEL1L-HRD1 (Ser658Pro) interfaces disrupt complex formation and impair ERAD activity. A newly identified disease-associated HRD1 variant (Ala91Asp), located in transmembrane helix 3, impairs HRD1 dimerization and substrate processing, underscoring the functional necessity of this interface and HRD1 dimerization. Finally, the structure also reveals two methionine-rich crevices flanking the HRD1 dimer, suggestive of substrate-conducting channels analogous to those in the ER membrane protein complex (EMC). These findings establish a structural framework for mammalian ERAD and elucidate how mutations destabilizing this machinery contribute to human disease.

**SUMMARY:** The dimeric structure of the human SEL1L-HRD1 ERAD core complex reveals key architectural and functional principles underlying the recognition and processing of misfolded proteins linked to human disease.

## INTRODUCTION

In eukaryotes, approximately 30% of newly synthesized proteins are directed to the endoplasmic reticulum (ER), where they fold and mature into their functional forms ^1,2^. Proteins that fail to fold properly are eliminated via ER-associated degradation (ERAD), a pathway that retrotranslocates misfolded proteins from the ER lumen to the cytosol for proteasomal degradation ^3–7^. The suppressor/enhancer of Lin-12-like protein 1-like (SEL1L) and the E3 ubiquitin ligase HMG-CoA reductase degradation protein 1 (HRD1) complex, homologous to the yeast Hrd3-Hrd1 pair, forms the core of the mammalian ERAD machinery and represents its most extensively characterized module ^8–12^. The discovery of pathogenic SEL1L and HRD1 variants in patients with ERAD-associated neurodevelopmental disorders with onset at infancy (ENDI syndrome) ^13,14^, along with the severe phenotypes observed in global and cell type-specific *Sel1L-* or *Hrd1*-knockout (KO) mice ^12,15,16^, underscores the critical role of ERAD in human health. However, the molecular mechanism by which these variants impair ERAD remains poorly understood, primarily due to the lack of structural information on the human complex.

Hrd3/SEL1L and Hrd1/HRD1 form the ERAD core, with HRD1 acting as the E3 ubiquitin ligase ^17^ and SEL1L stabilizing HRD1 ^18,19^ and promoting complex assembly ^20,21^. Additional cofactors include ER lectins Yos9/OS9 ^22–24^ and ERLEC1 (also known as XTP3B) ^25,26^, Der1/Derlin ^27–29^, and Usa1/HERP ^30^, which assist in substrate recognition, retrotranslocation, and ubiquitination. Previous cryo-electron microscopy (cryo-EM) studies of yeast ERAD revealed both a homo-dimeric Hrd3-Hrd1 complex ^31^ and monomeric Yos9-Hrd3-Hrd1 and Hrd3-Hrd1-Usa1-Derl subcomplexes ^32^. While the dimeric structure suggested a protein-conducting channel formed by the Hrd1 dimer, the dimer interface was closed, and the aqueous channel interior was instead exposed to the surrounding lipid layer ^31^. It was later proposed that this conformation to be a detergent artifact with low ubiquitination activity ^32^, favoring the monomeric model in yeast. In this monomeric model, Hrd1 and Der1 form complementary "half-channels" within a thinned ER membrane, while Hrd3 and Yos9 act as luminal substrate receptors ^32^. Whether similar assemblies exist in mammals remains unknown.

Notably, both size-exclusion chromatography and chemical-mediated crosslinking assays of endogenous SEL1L-HRD1 complex from human cells support HRD1 oligomerization ^33^ and dimerization ^34,35^. These findings raise the possibility that unlike the yeast complex, SEL1L-HRD1 dimerization is functionally important in mammals. In this study, we attempted to define the architecture of the mammalian ERAD core complex, which is essential for understanding the molecular basis of ERAD-related diseases.

## RESULTS

### The OS9-SEL1L-HRD1 complex forms the core of mammalian ERAD machinery

To first define the composition of the SEL1L-HRD1 ERAD core complex, we performed LC-MS/MS-based proteomic analyses of SEL1L and HRD1 interactomes in HEK293T cells following immunoprecipitation (IP) of either protein (Figure. S1A) ^36^. Each interactome was independently analyzed in triplicate using stringent filtering criteria, with *SEL1L* or *HRD1* KO cells as negative controls (Figure. S1A). We identified 46 high-confidence SEL1L interactors and 24 for HRD1 (Figures. S1B and S1C; Tables S1-S2). Cross-comparison revealed 10 shared high-confidence interactors (Figure. S1D; Table S3), with OS9, ERLEC1, SEL1L, HRD1, and FAM8A1 emerging as the most prominent components (Figure. 1A). OS9 and ERLEC1 are ER lectins with similar domain architectures and redundant roles in HRD1-mediated ERAD ^26,37^. Consistent with previous findings ^38^, FAM8A1 deletion had no effect on ERAD function or SEL1L-HRD1 protein levels (Figure. S1E).

**Figure 1.**
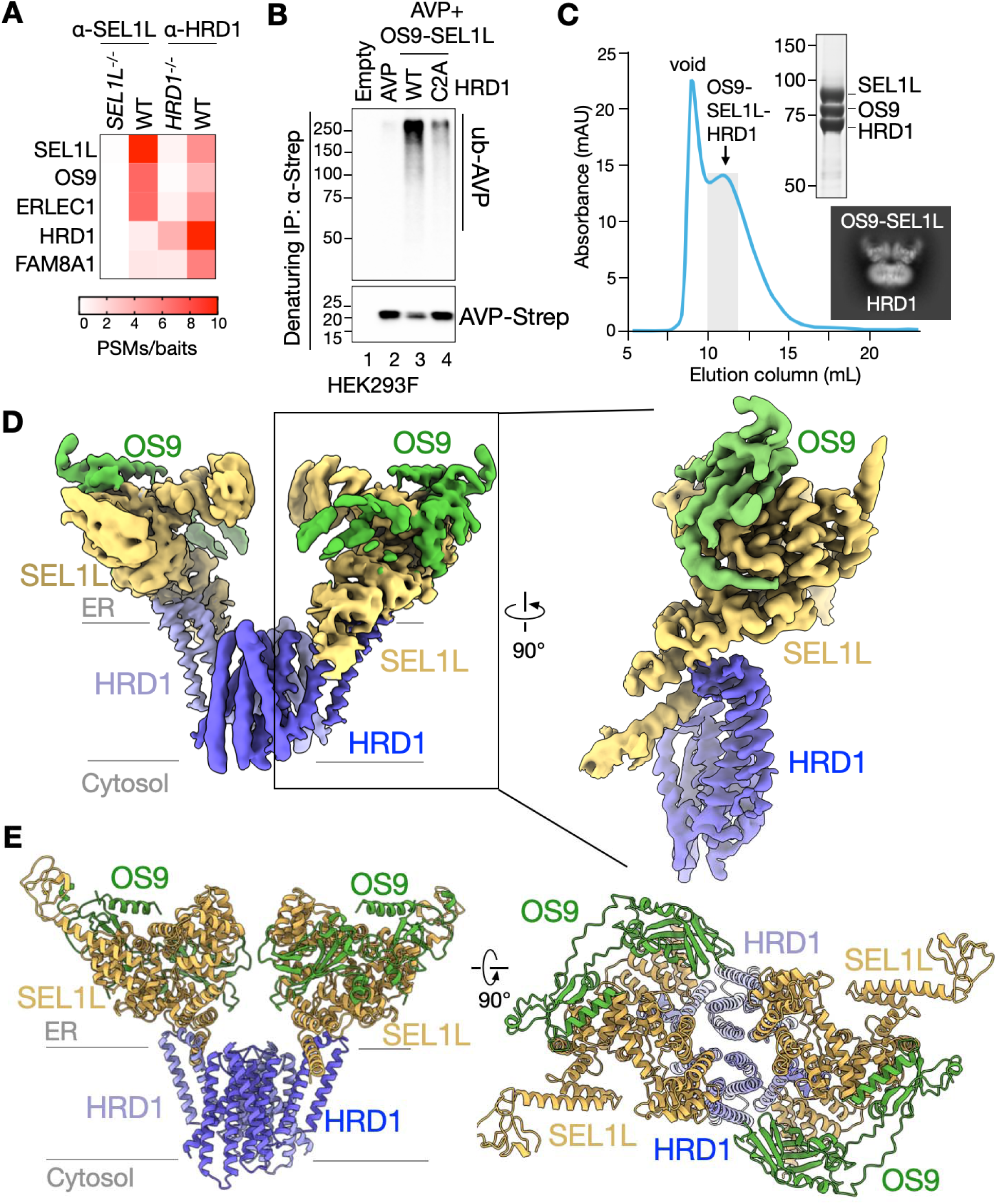
Cryo-EM structure of dimeric OS9-SEL1L-HRD1 ERAD core complex. (**A**) Heatmap showing the top 5 hits of the overlapping 10 hits of SEL1L- and HRD1-interacting proteins from immunoprecipitation-Mass Spec (IP-MS) experiments (Fig. S1D and Table S3). Plotted values are peptide-spectrum matches (PSMs) normalized to their baits from three independent experiments. (**B**) Denaturing immunoprecipitation of Strep-agarose in HEK293F cells expressing the ERAD substrate proAVP(G57S)-Strep, OS9, SEL1L, and HRD1-WT-FLAG or the ligase-dead HRD1 mutant C2A (C291A/C294A)-FLAG, with 10 μM MG132 for 2 hours, to measure proAVP(G57S) ubiquitination (two independent repeats). Input is shown in Fig. S2C. (**C**) Size exclusion chromatography (SEC) of OS9-SEL1L-HRD1 complex. FLAG-tagged HRD1, untagged SEL1L and untagged OS9 are expressed in HEK293F cells. The OS9-SEL1L-HRD1 ERAD complex was purified with FLAG-agarose and the elution was subjected to gel filtration on a Superose 6 10/300GL increase column. The peak fractions (10-11 mL, shaded) were concentrated for cryo-EM sample preparation. The right panels show a Coomassie-stained SDS-PAGE gel of the concentrated OS9-SEL1L-HRD1 ERAD complex for cryo-EM sample preparation, and representative 2D class averages of picked particles. (**D**) Cryo-EM map of the dimeric OS9-SEL1L-HRD1 complex. Inset showing the locally refined OS9-SEL1L map stitched with HRD1, as a monomer, for visualization. (**E**) Cartoon model of the OS9-SEL1L-HRD1 dimer rotated to show the top view of the structure.

To further define the core components of the SEL1L-HRD1 complex, we co-expressed key factors (HRD1, SEL1L, OS9, HERP1, UBE2J1, and DERL2) in HEK293F cells, followed by HRD1 pull-down assays. The OS9-SEL1L-HRD1 subcomplex was robustly co-purified under various detergent conditions, consistently maintaining a stable stoichiometric ratio, except in Triton X-100 and NP-40, which disrupted complex integrity (Figures. S2A and S2B). Co-expression of OS9, SEL1L, and HRD1 in HEK293F cells with a model HRD1-ERAD substrate, pro-arginine vasopressin (proAVP) harboring the Gly57Ser disease mutation prone to misfolding ^39^, resulted in substrate ubiquitination in cells (Figures. 1B and S2C). This activity was abolished upon expression of a ligase-dead HRD1 mutant (C291A/C294A; C2A) ^40^ (Figures. 1B and S2C). Taken together, these findings establish the OS9-SEL1L-HRD1 complex as the functional core of the mammalian ERAD machinery.

### OS9-SEL1L-HRD1 ERAD core complex forms a dimer

To determine the molecular organization of the OS9-SEL1L-HRD1 core complex, we co-expressed full length OS9, SEL1L and FLAG-tagged HRD1 in HEK293F cells. Following membrane solubilization in lauryl maltose neopentyl glycol (LMNG), the complex was purified by FLAG-based immunoprecipitation and size-exclusion chromatography. This procedure yielded a single, well-defined peak containing all three components – OS9, SEL1L and HRD1 (Figures. 1C and S2D). Peak fractions were collected, concentrated and immediately prepared for sample freezing and subsequent structural determination by cryo-EM single-particle analysis (Figures. S3A and S3B).

Initial 2D image analysis revealed the presence of a homogenous set of particles, most of which display visible C2 symmetry (Figure. S3A). Subsequent 3D reconstruction with the selected particles revealed that the OS9-SEL1L-HRD1 complex adopts a dimeric architecture, with an overall resolution of 3.6 Å (Figures. S3A and S3B; Table S4). We then performed a focused 3D refinement using a mask around the visible regions of OS9 and SEL1L in the structure (Figure. S3B). This process led to improvements in local resolutions to 3.3 Å for the OS9-SEL1L interface (Figures. 1D and S3B). Model building into the EM density maps was guided using AlphaFold3-predicted structures ^41^, followed by manual adjustments based on visible side chain densities for SEL1L-OS9 and the transmembrane helices of HRD1. This enabled accurate modeling of the three core components: OS9, SEL1L, and HRD1 (Figures. 1D-1E and S4A-S4E). The final 3D model captured all major domains of the core ERAD complex: the eight transmembrane helices of HRD1 (residues 1-265), the luminal domain of SEL1L (residues 119-725), and two conserved domains of OS9 – the mannose-6-phosphate receptor homology (MRH) domain (residues 88-236) and a structured region comprising four β-sheets and an α-helix (termed “4β”, residues 28-87, 237-259, 633-652) (Figures. 1D-1E and S4B-S4E). Regions not resolved in the map included the transmembrane domain of SEL1L, the cytosolic regions of both SEL1L and HRD1 (including the E3 ligase RING domain), and the remaining flexible portions of OS9’s luminal domain (Figure. S4A).

### SEL1L disease variants disrupt the OS9-SEL1L-HRD1 core complex

SEL1L, which contains multiple Sel-like tetratricopeptide repeats (SLR) and amphipathic helix (APH) (Figures. S4A, S4B and S4D), serves as a central scaffold in the ERAD complex by linking the ER lectin OS9 to the E3 ligase HRD1 ^21,25,36^. To establish the OS9-SEL1L-HRD1 assembly as the functional core of mammalian ERAD *in vivo*, we examined two disease-causing SEL1L variants, G585D and S658P, previously shown to impair ERAD function ^13,36^. Our cryo-EM map of the OS9-SEL1L subcomplex revealed a crab claw-like configuration with a central pore (asterisk, Figure. 2A), suggestive of a conduit for OS9-bound substrate delivery to HRD1. SEL1L-G585 maps to the SEL1L-OS9 interface (Figures. 2B, 2C and S5A), while SEL1L-S658 lies near the SEL1L-HRD1 interface (Figures. 2D, 2E and S5A). In *SEL1L^-/-^* HEK293T cells, reconstitution with the G585D mutant selectively abrogated OS9 binding (lane 3, Figure. 2F), while S658P disrupted HRD1 interaction (lane 4, Figure. 2F). The double mutant, G585D/S658P, abolished both interactions, effectively dismantling the core complex (lane 5, Figure. 2F). Importantly, neither mutation affected binding to the E2 conjugating enzyme UBE2J1 (lane 2-5, Figure. 2F), indicating interaction specificity. These data establish the OS9-SEL1L-HRD1 complex as the structural and functional core of the mammalian ERAD machinery.

**Figure 2.**
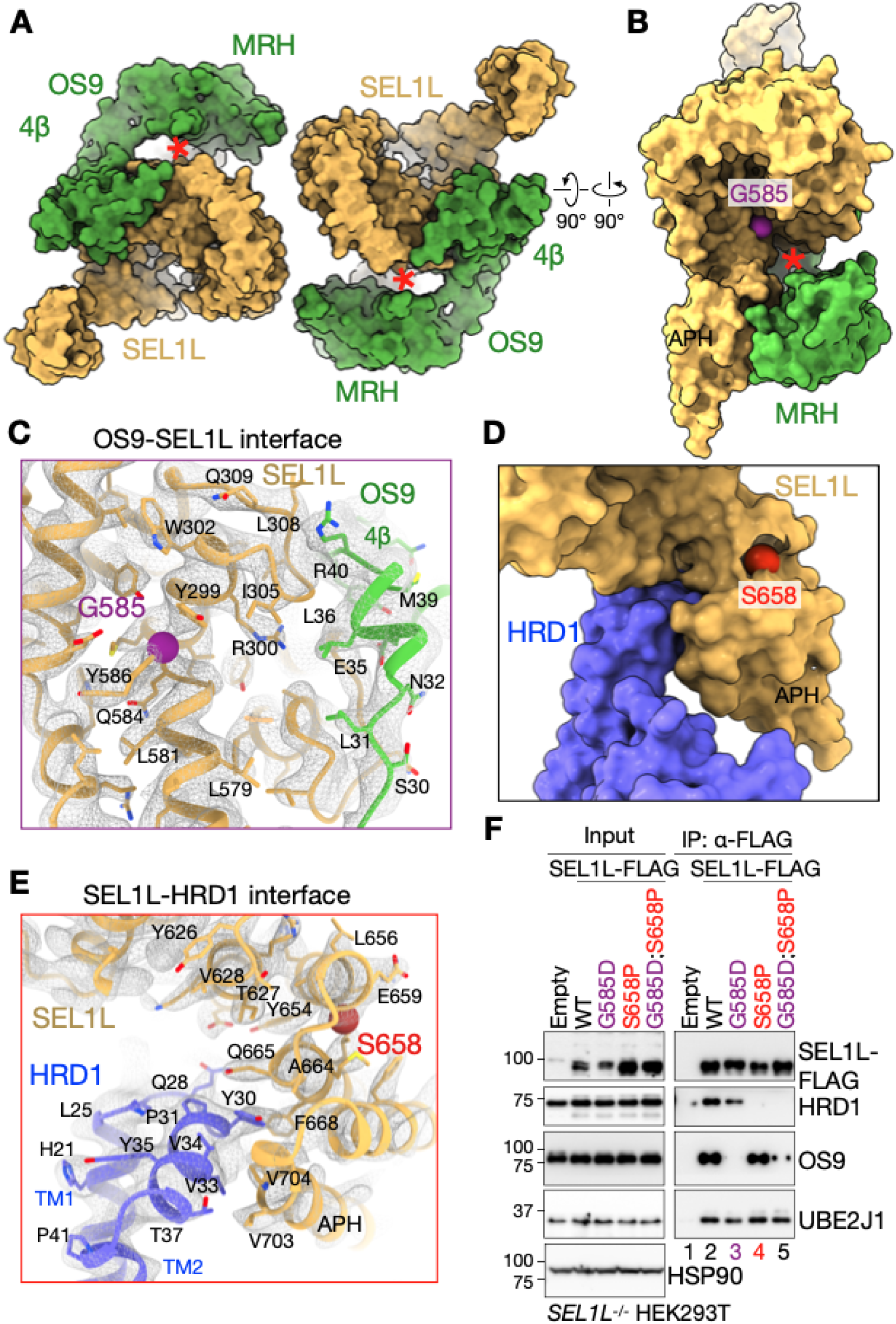
SEL1L disease variants disrupt OS9-SEL1L and SEL1L-HRD1 interaction. (**A-B**) View from ER lumen (**A**) or the side (**B**) with space-filling model for OS9-SEL1L complex. Red asterisk indicates the opening formed by SEL1L-OS9 in substrate-binding crab “claw”-like configuration. The position of SEL1L-G585 residue (purple sphere) shown in (**B**). 4β, the four β-sheet domain of OS9. MRH, mannose-6-phosphate receptor homology domain of OS9. APH, amphipathic helix of SEL1L. (**C**) Interface of OS9-SEL1L interaction with SEL1L-G585 residue shown as a purple sphere, with the cryo-EM map shown as mesh. (**D**) Close-up view with space-filling model for SEL1L-HRD1 showing the SEL1L-HRD1 interaction interface, with SEL1L-S658 residue shown as a red sphere. APH, amphipathic helix of SEL1L. (**E**) Interface of SEL1L-HRD1 interaction with SEL1L-S658 residue shown as a red sphere, with the cryo-EM map shown as mesh. APH, amphipathic helix of SEL1L. (**F**) Immunoprecipitation of FLAG-agarose in *SEL1L*^-/-^ HEK293T cells expressing indicated SEL1L variants to examine their interaction with HRD1, OS9 and E2 UBE2J1 (three independent repeats).

### *In vivo* formation of OS9-SEL1L dimers

Unlike the monomeric OS9-Hrd3 structure in yeast ^32^, our cryo-EM structure revealed that OS9-SEL1L protomer adopts a closed dimer interface (Figure. 3A). Within this interface, OS9 residue D135 lies in close proximity to SEL1L residues D439 and M440 from the opposing protomer (Figures. 3B and 3C). To test whether this dimerization occurs *in vivo* – and is not an artifact of detergent-based purification – we performed native disulfide crosslinking assays on ER membrane-associated complex ^42^. To prevent disulfide bond formation during lysis, cells were lysed directly in non-reducing SDS buffer rather than conventional Triton X-100 or NP-40 lysis buffers. In the oxidizing environment of the ER lumen ^43^, such spatial proximity enables disulfide bond formation between engineered cysteines ^42^. We introduced D135C into OS9 and D439C or M440C into SEL1L, co-expressed with HRD1-FLAG. SEL1L-WT and the more distal N442C mutant served as negative controls (Figures. 3B-3D). Disulfide-linked complexes formed between OS9 D135C and SEL1L D439C or M440C but was substantially reduced with N442C and absent with WT (lanes 7-8 vs. 9-10, Figure. 3D). Coomassie blue staining following HRD1-FLAG IP confirmed OS9-SEL1L complex formation under native conditions (lane 1 vs. 2, Figure. S6A). In all these assays, these disulfide-linked complexes remained stable under denaturing conditions, but were disrupted by reducing agents (lane 2-3 vs. 7-8, Figure. 3D; lane 1 vs. 3, Figure. S6A), supporting *in vivo* disulfide-mediated dimerization of the OS9-SEL1L complex.

**Figure 3.**
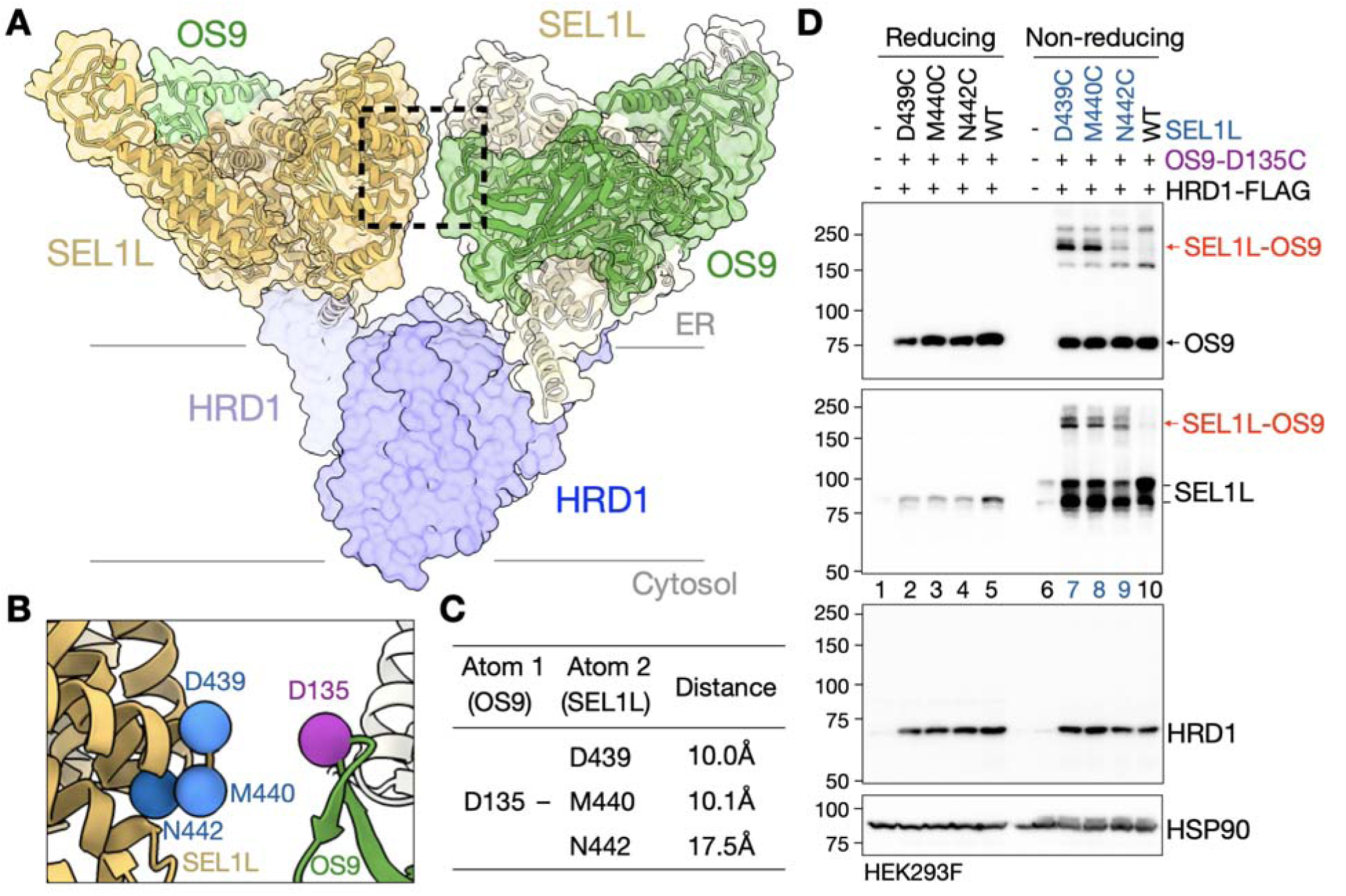
Homodimeric OS9-SEL1L complex forms *in vivo*. (**A-B**) OS9-SEL1L-HRD1 model shown as a transparent surface. Zoomed-in view of the dashed box shown in (**B**). Key residues (SEL1L-D439, M440, and OS9-D135) at the interface, and negative control SEL1L-N442 are highlighted. (**C**) Distance between OS9-D135 and SEL1L-D439, M440 and N442. (**D**) Reducing and non-reducing SDS-PAGE and Western blot analysis of the native disulfide crosslinking assays in HEK293F cells expressing HRD1-FLAG, SEL1L WT and mutants (D439C, M440C, N442C), and OS9-D135C mutations (two independent repeats).

### *In vivo* formation of HRD1 dimers

Our cryo-EM structure revealed that HRD1 dimerizes via interactions at transmembrane helix 3 (TM3) (Figure. 4A). Residues T93, F95 and R96 were positioned facing each other or were within close proximity at the dimer interface, while V94 and D97 were located more peripherally and oriented away from the interface (Figures. 4B and S7A). Several of these residues are evolutionarily conserved (Figure. 4C). To probe the dimerization interface, we introduced cysteine substitutions at these positions. Native disulfide crosslinking assay showed that T93C, F95C, and R96C efficiently formed disulfide-linked HRD1 dimers, with approximately 40% dimerization, which were abolished by reducing agents (lanes 3, 5, 6, Figure. 4D), indicating a physiologically relevant dimeric interface in the ER membrane. In contrast, crosslinking was markedly reduced in cells expressing V94C and D97C (lanes 4 and 7, Figure. 4D). To further confirm endogenous HRD1 dimerization *in vivo*, we treated HEK293T cells with the membrane-permeable crosslinker dithiobis-succinimidyl propionate (DSP), previously used to study HRD1 dimer complexes ^34,35^. Consistent with our structural and biochemical findings, HRD1 dimers were readily detected in WT cells but were markedly reduced in SEL1L-deficient cells (Figure. S7B). In both crosslinking assays, the dimeric complexes were sensitive to reducing agents, confirming the presence of disulfide/DSP-linked interactions (Figures. 4D and S7B). Taken together, these findings demonstrate that the OS9-SEL1L-HRD1 complex forms a stable, stoichiometric dimer *in vivo*.

**Figure 4.**
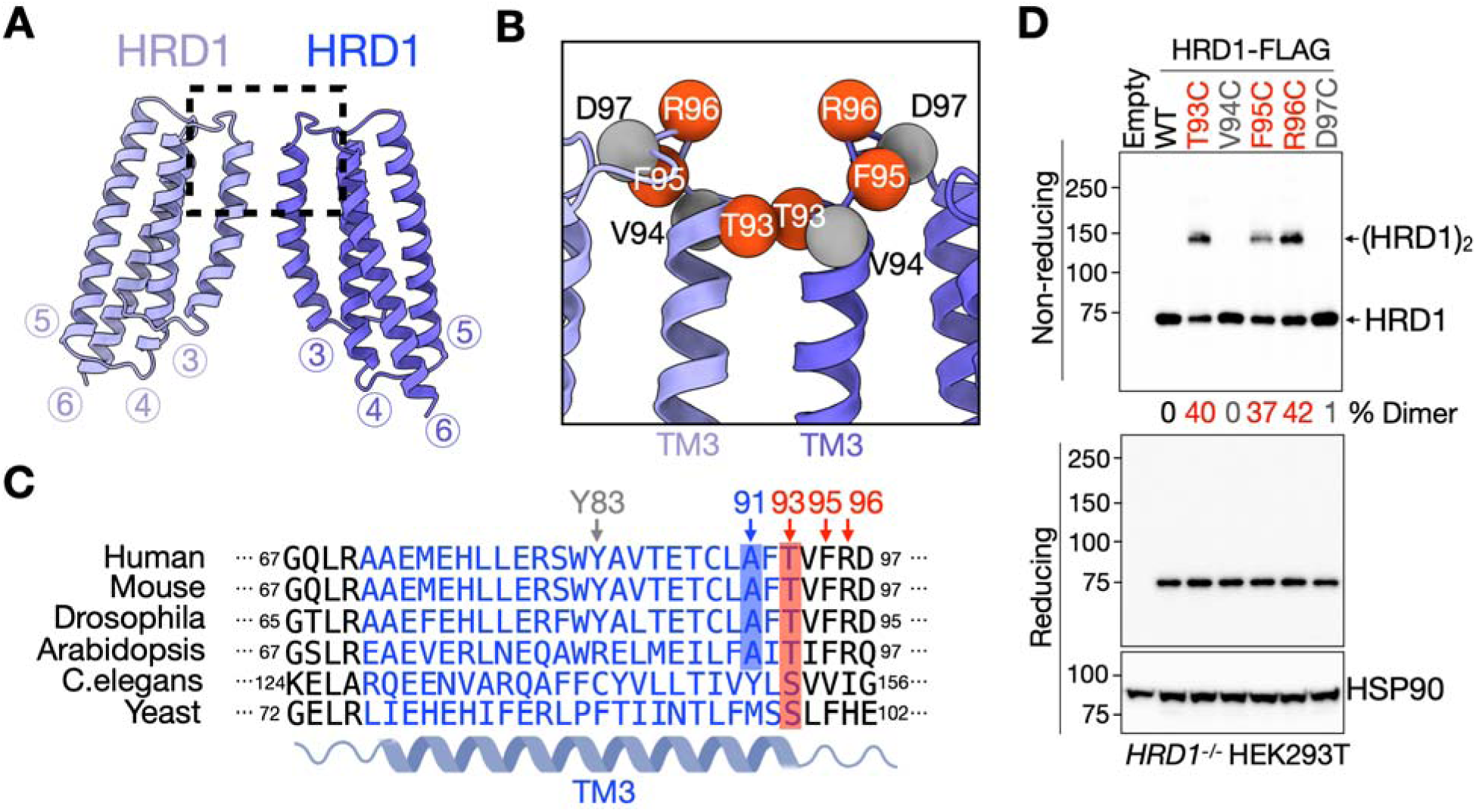
HRD1 forms a homodimer *in vivo*. (**A-B**) Overview of the HRD1 homodimer. The numbers below the model refer to the TMs of HRD1. The dashed box indicates the interface of HRD1 dimer. Zoomed-in view of the dashed box shown in (**B**). Key residues (HRD1-T93, F95 and R96) in HRD1 TM3 or the loop between TM3 and 4 are highlighted. (**C**) Sequence alignment of TM3 of HRD1 across various model organisms, identifying the conservation of HRD1-T93, F95 and R96 among various organisms. (**D**) Reducing and non-reducing SDS-PAGE and Western blot analysis of the native disulfide crosslinking assays in *HRD1^-/-^* HEK293T cells expressing the indicated HRD1 variants. The percentage of the dimer HRD1 in total HRD1 is shown below the gel (Three independent repeats).

### TM3-mediated HRD1 dimerization is required for function

In yeast, Hrd1 dimers have been proposed to be inactive ^32^. We next delineated the functional relevance of HRD1 dimers in mammalian cells. Based on the cryo-EM structure map, HRD1 homodimerization is stabilized by a polar interaction involving T93 and a π-stacking interaction mediated by Y83 within TM3 of opposing HRD1 protomers (Figures. 5A and S8A). While Y83 is not conserved and appears to contribute less to the interface based on the EM density, T93 is highly conversed from yeast to humans and is likely the dominant stabilizing interaction (Figures. 4C and S8A). To assess the role of T93, we introduced T93A and T93F mutants to disrupt the polar interaction and examined dimerization using R96C-mediated native disulfide crosslinking and DSP-based chemical crosslinking. Both mutations abolished disulfide-linked dimer formation (lane 4-5 vs. 3, Figure. 5B) and reduced chemically crosslinked dimers by ∼10-fold (lanes 3-4 vs. 2, Figure. 5C). These data demonstrate that the polar interaction at T93 is critical for TM3-mediated HRD1 dimerization.

**Figure 5.**
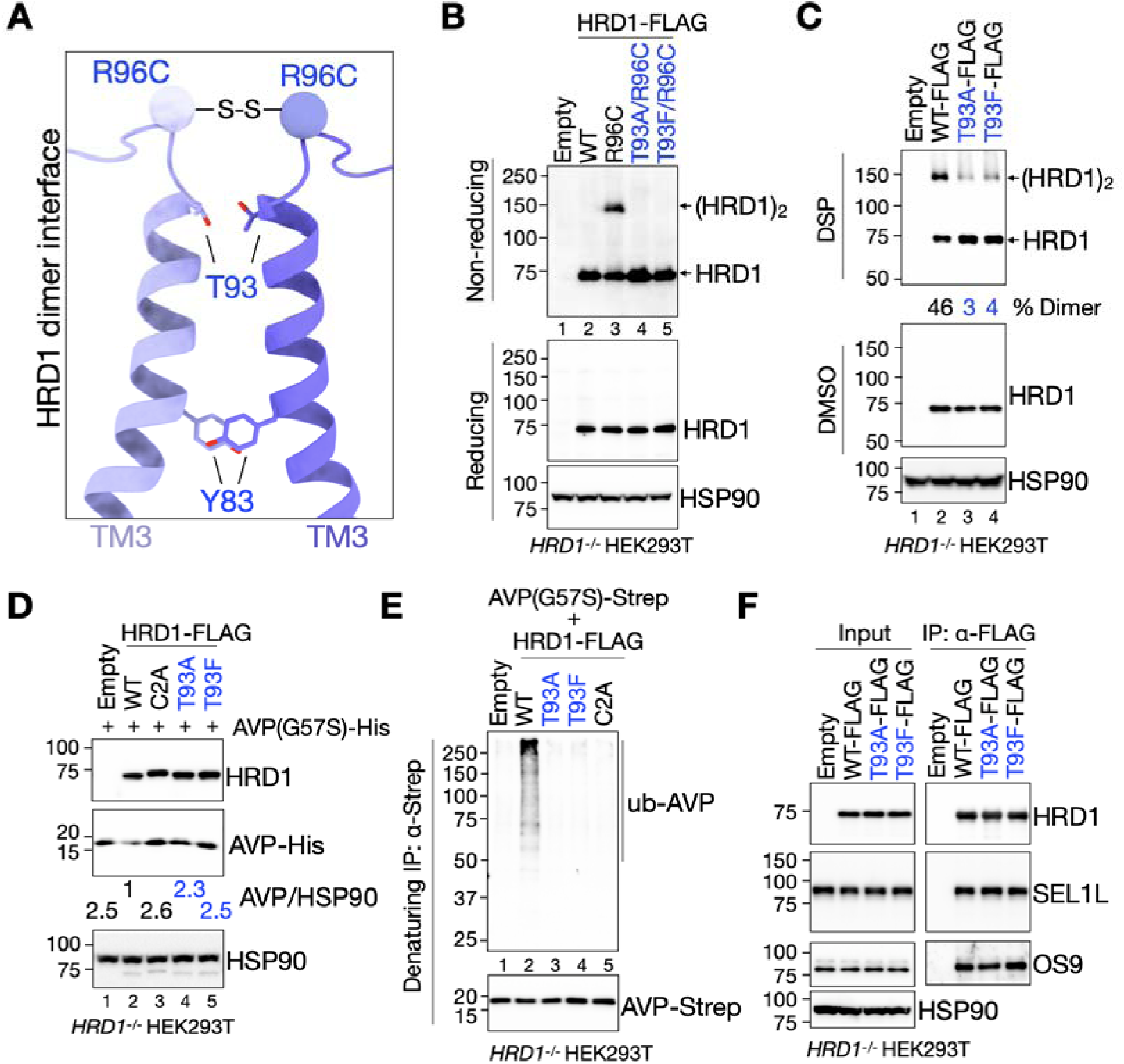
Interface of HRD1 dimer on TM3 is required for ERAD function. (**A**) Side view of HRD1 TM3 at the dimer interface. The density maps showing the polar interaction of T93-T93 and Y83-Y83 are shown in Fig. S8B. S-S indicates the disulfide bond formed by HRD1-R96C. (**B**) Reducing and non-reducing SDS-PAGE and Western blot analysis of the native disulfide crosslinking assays in *HRD1^-/-^* HEK293T cells expressing the indicated HRD1 mutants (three independent repeats). (**C**) Reducing and non-reducing SDS-PAGE and Western blot analysis of DSP-mediated crosslinking in *HRD1^-/-^* HEK293T cells expressing indicated HRD1 mutants, following 2-hr 2 mM DSP treatment. The percentage of the dimer HRD1 in total HRD1 is shown below the gel (two independent repeats). (**D**) Western blot analysis of ERAD substrate proAVP(G57S) levels in *HRD1^-/-^* HEK293T cells expressing the indicated HRD1 mutants, with quantitation of proAVP(G57S) levels shown below the gel normalized to cells expressing WT HRD1 (three independent repeats). (**E**) Denaturing IP of Strep-agarose in *HRD1^-/-^* HEK293T cells expressing proAVP(G57S)-Strep and indicated HRD1 mutants, treated with 10 μM MG132 for 2 hours, to measure proAVP(G57S) ubiquitination (two independent repeats). (**F**) Immunoprecipitation of FLAG-agarose in *HRD1*^-/-^ HEK293T cells expressing indicated HRD1 mutants to examine their interaction with SEL1L and OS9 (two independent repeats).

We next examined whether HRD1 dimerization is required for ERAD function. Expression of the model substrate proAVP(G57S) in *HRD1^-/-^*HEK293T cells reconstituted with either HRD1 T93A or T93F mutants led to substrate accumulation (lanes 3-4 vs. 2, Figure. 5D) and impaired ubiquitination (lanes 3-4 vs. 2, Figure. 5E), similar to the catalytically inactive HRD1-C2A mutant (lane 3, Figure. 5D; lane 5, Figure. 5E). These defects were not due to disruption of the core complex as both T93 mutants retained binding to OS9 and SEL1L (Figure. 5F), suggesting that HRD1 dimerization is dispensable for assembly of the monomeric OS9-SEL1L-HRD1 complex. Thus, the T93-mediated polar interaction is critical for HRD1 dimerization and ERAD function, independent of OS9 and SEL1L binding.

### A disease-associated *HRD1* variant disrupts dimerization and impairs ERAD function

We recently identified a novel, potential disease-causing HRD1 variant, *A91D*, in a one-year-old boy with congenital heart defects and early-onset pulmonary dysfunction, born to consanguineous parents (Lifera Omics Database). A91 is conserved from humans to Arabidopsis but not in yeast (Figure. 4C), and our cryo-EM structure located this residue within TM3 of HRD1 at the dimer interface (Figure. 6A). Expression of HRD1 A91D in *HRD1^-/-^* HEK239T cells severely impaired degradation of the misfolded substrate proAVP(G57S), leading to substrate accumulation (lane 4 vs. 3, Figure. 6B) and ∼70%-reduction in ubiquitination (lane 5 vs. 4, Figure. 6C) – indicative of ERAD dysfunction. Native disulfide crosslinking using the R96C reporter, as well as DSP-based chemical crosslinking, revealed a > 60% reduction in dimer formation by the A91D mutant (Figures. 6D and 6E). Importantly, the A91D variant retained normal binding to OS9 and SEL1L (Figure. 6F), indicating that the defect is specific to HRD1 dimerization rather than disruption of the core complex. These findings identify HRD1 A91D as a pathogenic variant that selectively disrupts HRD1 dimerization and ERAD activity, directly linking dimer interface integrity to human disease pathogenesis.

**Figure 6.**
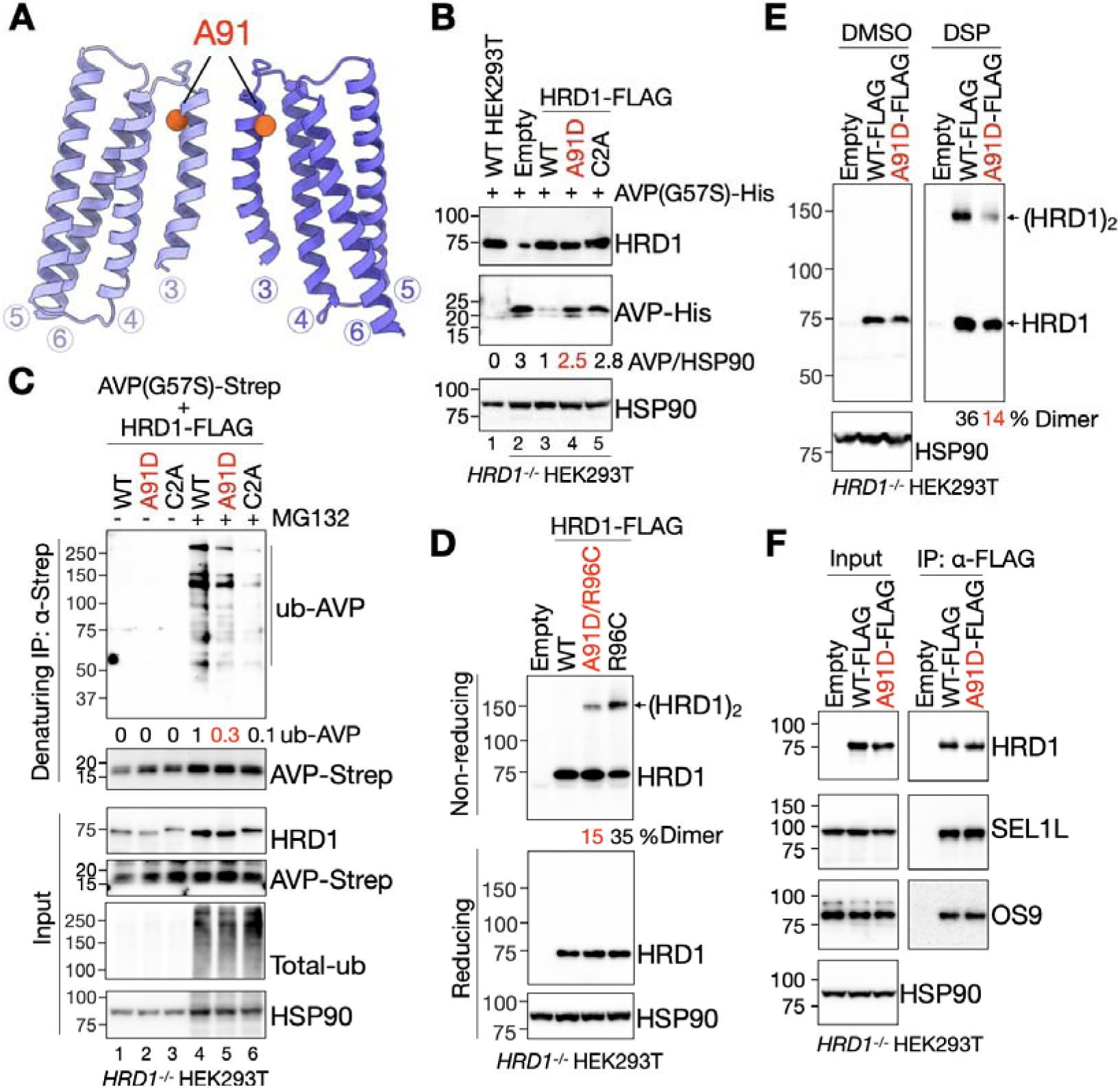
*HRD1^A91D^* disease variant abolishes HRD1 dimerization and function. (**A**) HRD1 dimer model indicating the position of A91 as orange spheres. (**B**) Western blot analysis of proAVP(G57S) protein levels in *HRD1^-/-^* HEK293T cells expressing the indicated HRD1 mutants, with quantitation of proAVP(G57S) levels shown below the gel normalized to cells expressing WT HRD1 (three independent repeats). (**C**) Denaturing IP of Strep-agarose in *HRD1^-/-^* HEK293T cells expressing proAVP(G57S)-Strep and indicated HRD1 mutants, with or without 10 μM MG132 for 2 hr, to measure proAVP(G57S) ubiquitination (two independent repeats). (**D**) Reducing and non-reducing SDS-PAGE and Western blot analysis of the native disulfide crosslinking assays in *HRD1^-/-^* HEK293T cells expressing the indicated HRD1 mutants (two independent repeats). (**E**) Reducing and non-reducing SDS-PAGE and Western blot analysis of DSP-mediated crosslinking in *HRD1^-/-^* HEK293T cells expressing indicated HRD1 mutants, wi following 2-hr 2 mM DSP treatment. The percentage of the dimer HRD1 in total HRD1 is shown below the gel (two independent repeats). (**F**) Immunoprecipitation of FLAG-agarose in *HRD1*^-/-^ HEK293T cells expressing HRD1-WT and A91D variant to examine their interaction with SEL1L and OS9 (two independent repeats).

### A methionine-lined channels in the HRD1 dimer suggests a role in substrate translocation

Lastly, we examined whether the HRD1 dimer harbors a potential protein-conducting channel. Our cryo-EM structure revealed a dimeric HRD1 architecture that differs markedly from the previously reported yeast Hrd1-Hrd3 dimer ^31^ and Hrd1-Der1 complex ^32^ (Figures. 7A-7C and S9A-S9C). We identified a putative translocation channel formed between TM1-4 of one HRD1 protomer and TM8 of the other (asterisks, Figures. 7A and S9A) – a configuration that was not observed in the yeast Hrd1-Hrd3 dimer (Figures. 7B and S9B) and also distinct from the Hrd1-Der1 complex, where paired half-channels are formed by TM2-4 of Hrd1 and TM1-2, TM5 of Der1 (asterisks, Figures. 7C and S9C).

**Figure 7.**
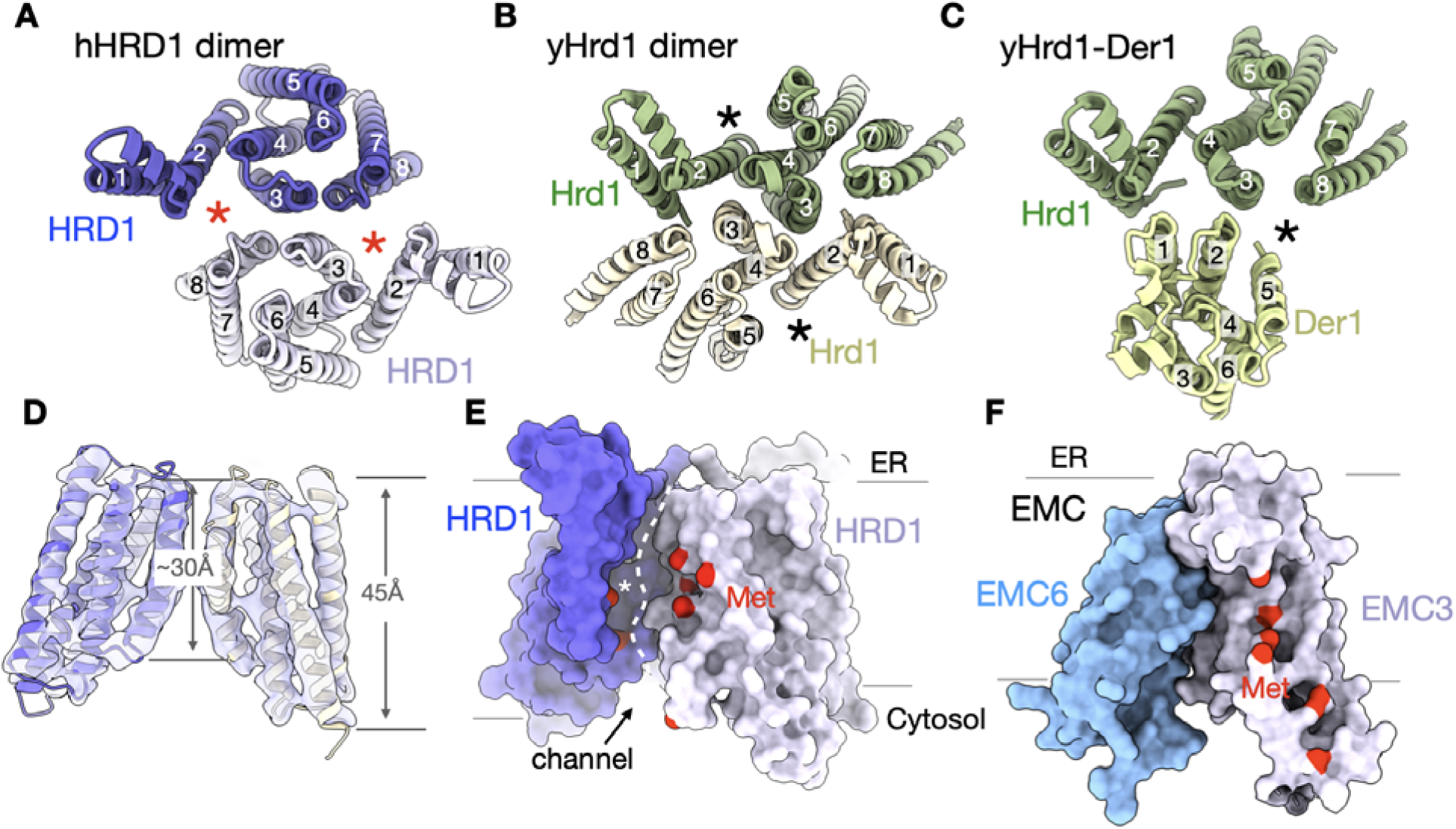
HRD1 dimer forms putative substrate-conductive channel lined with methionine-rich bristles within a locally thinned membrane region. (**A-C**) Comparison of the human HRD1 dimer (hHRD1) (**A**), yeast Hrd1 dimer (yHrd1) (PDB code: 5V6P) (**B**), and yeast Hrd1-Der1 structures (PDB code: 6vjz) (**C**). Asterisks indicate the putative substrate retrotranslocation channels. (**D**) Side view of the HRD1 model. The cryo-EM map filtered to 4Å is shown as a transparent surface. The distances of the HRD1 interface (TM3) and the peripheral region are shown in red. (**E-F**) Surface model of the membrane-embedded region of HRD1 dimer and human EMC3-6 subcomplex (**F**). Methionine (Met) residues in the putative substrate retrotranslocation channel are highlighted in red, showing methionine-conducted bristles. EMC model is generated by AlphaFold-3 as methionine-conducted bristles are invisible in the EMC structure. Asterisk and the dashed line in (**E**) indicate the putative substrate retrotranslocation channel.

Analogous to the inward-bent helices in the Hrd1-Der1 complex that create a substrate-binding cavity ^32^, the HRD1 dimers appears to induce membrane thinning or bending at the dimer interface, forming the cavity observed in the micelle EM density (Figures. 7D and S9D). Strikingly, this channel is enriched in methionine residues (red, Figure. 7E), reminiscent of the “methionine bristles” that line the translocation pores of the ER membrane protein complex (EMC) and bacterial YidC insertase ^44,45^, as well as the M domain of the signal recognition particle ^46,47^ (Figures. 7F and S9E). These findings reveal a previously unrecognized methionine-lined channel within the HRD1 dimer, suggesting a possible role in substrate engagement or translocation during ERAD.

## DISCUSSION

To define the molecular organization of the human ERAD machinery, we performed an unbiased IP-MS screen to define its core components, followed by single-particle cryo-EM to determine the structure of the OS9-SEL1L-HRD1 complex. The structure reveals that this core complex forms a dimer, with HRD1 assembling as a symmetric homodimer (Figure. 8A). The two HRD1 protomers are arranged in parallel, primarily stabilized by interactions between TM3 of opposing protomers. Functional validation using structure-guided mutagenesis – including the disease-associated HRD1 variant A91D – demonstrated that perturbing this dimer interface impairs complex formation and ERAD activity (Figure. 8B). This dimeric configuration not only reinforces the transmembrane assembly but also gives rise to a central cavity that may facilitate substrate engagement and retro-translocation.

**Figure 8.**
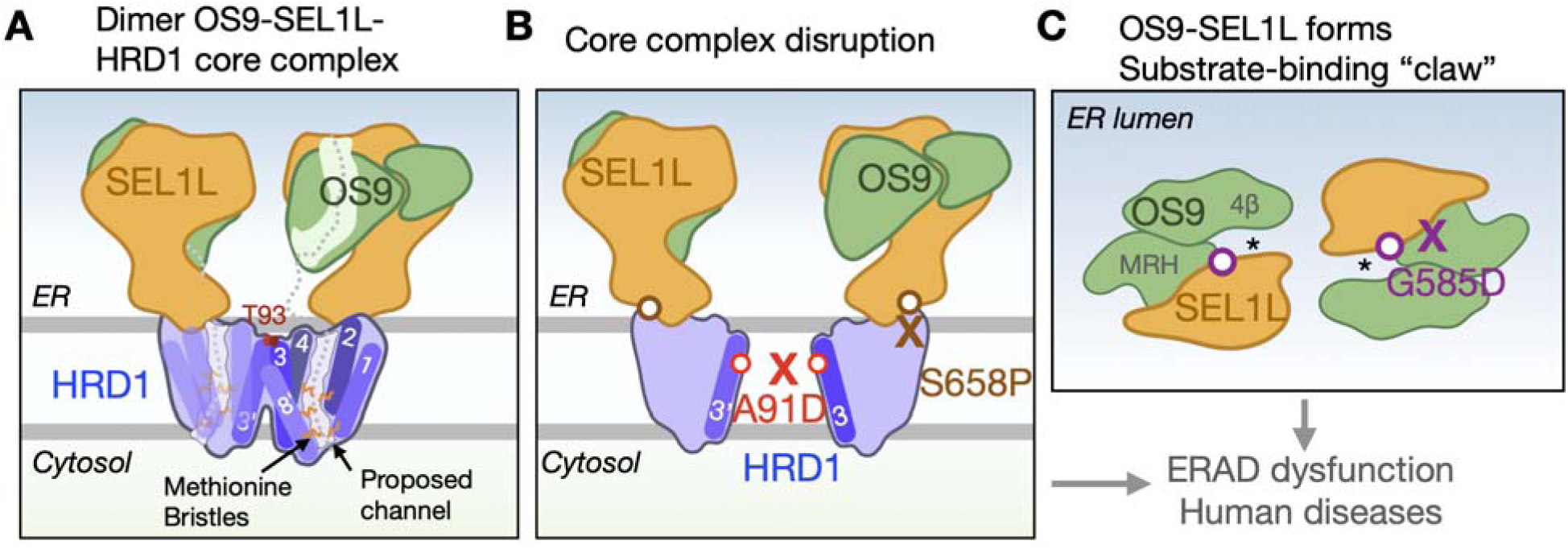
Schematic of the interaction interfaces and putative substrate binding regions in the structure of OS9-SEL1L-HRD1. (**A**) The OS9-SEL1L-HRD1 ERAD core complex forms a homodimer *in vivo*, with HRD1 dimerization dependent on polar interactions between two T93 residues - critical for ERAD function. In the ER lumen, the MRH domain of OS9 and SEL1L together form a substrate-binding “claw”-like configuration, while TM1-4 of one HRD1 and TM8 of the opposing protomer create methionine bristles as a putative ERAD substrate retrotranslocation channel. (**B**) The HRD1 disease variant A91D disrupts dimerization, while the SEL1L variant S658P impairs the SEL1L-HRD1 interaction - both compromising ERAD complex formation and function. (**C**) ER-lumen view of the OS9-SEL1L complex highlights the substrate-binding crab claw-like configuration. SEL1L disease variant G585D disrupts the SEL1L-OS9 interaction. All three disease mutants attenuate ERAD function via distinct mechanisms.

Our combined proteomic screen and structural analysis identify OS9, SEL1L, and HRD1 as the minimal core components of the human ERAD complex. Within this architecture, SEL1L acts as a scaffold, bridging luminal lectin OS9 and membrane-embedded HRD1. This configuration likely enables the recognition of misfolded proteins by OS9 and their directed transfer to HRD1 (Figure. 8A). Notably, Derlin – an essential translocation factor in yeast ERAD ^32^ – was not detected in our core complex, consistent with prior models proposing that Derlin functions downstream of HRD1 in mammals to recruit the cytosolic ATPase p97 ^27–29,48^. The absence of Derlin in this complex highlights an evolutionary divergence in ERAD mechanisms and raises the possibility that HRD1 itself functions as the principal protein-conducting channel in mammalian cells.

Based on these structural insights, we propose a model in which substrate capture occurs through a crab claw-like configuration formed by SEL1L and OS9, directing polypeptides toward a methionine-rich cavity at the HRD1 dimer interface (Figure. 8C). Each protomer contributes TM1-3 and TM8 to form two methionine-rich, hydrophobic grooves flanking the dimer axis (Figure. 8A). Methionine residues, with their flexible and nonpolar side chains, are well suited to form a dynamic, low-energy interface for transient substrate interactions during translocation. This configuration may allow HRD1 to accommodate the diverse structural and biochemical properties of misfolded substrates. Together, these findings define the molecular architecture of the mammalian ERAD core complex and provide a mechanistic framework for understanding substrate recognition and retro-translocation – essential processes disrupted in human disease.

## AUTHOR CONTRIBUTION

This study was conceived by L.L.L, A.J. and L.Q. L.L.L. purified the ERAD core complex and collected cryo-EM data. E.M. processed cryo-EM data and built the atomic model. L.L.L designed, performed all the biochemical experiments. L.E.Z. conducted the site directed point mutagenesis and plasmid preps. L.L.L wrote the manuscript. All authors contributed to data analysis and final version of the manuscript.

## ACKNOWLEDGEMENTS

We thank the Qi, Jomaa, Sun, Arvan Labs, Drs. David Castle, Qinli Hu (UT Southwestern Medical Center), Maciej Gluc, Haoran Yuan, Ruoya Ho, and Huan Bao for technical support and insightful discussions; Drs. Michael Purdy and David Cooper for assisting with the cryo-EM data collection and computational support; Dr. Chih-Chi Andrew Hu (Houston Methodist Hospital) for reagents; Riley Loyd for initiating cryo-EM data processing; and Drs. Fowzan Alkuraya and Khadijah Bakur for providing the information on the HRD1 A91D variant from the Lifera Omics Database. Cryo-EM data collection was conducted at the Molecular Electron Microscopy Core Facility (RRID:SCR_019031) at the University of Virginia (UVA) School of Medicine (NIH G20-RR31199). We thank the Proteomics Resource Facility at University of Michigan Medical School for assisting proteomic assays.

## Data and materials availability

The materials and reagents used are either commercially available or available upon the request. Proteomic data has been deposited into a public database PRIDE (PXD043674 and PXD041882). The cryo-EM map and the corresponding atomic model for the ERAD complex structure have been deposited under the accession codes EMD-70448 (consensus map), EMD-70452 (locally refined map) and PDB-9OG0. All data related to this study are available in the main text or the supplementary material.

## REFERENCE

1 Kanapin, A. et al. Mouse proteome analysis. Genome Res 13, 1335–1344, doi:10.1101/gr.978703 (2003).

2 Benham, A. M. Protein secretion and the endoplasmic reticulum. Cold Spring Harbor perspectives in biology 4, a012872, doi:10.1101/cshperspect.a012872 (2012).

3 Lippincott-Schwartz, J., Bonifacino, J. S., Yuan, L. C. & Klausner, R. D. Degradation from the endoplasmic reticulum: disposing of newly synthesized proteins. Cell 54, 209–220, doi:10.1016/0092-8674(88)90553-3 (1988).

4 Sommer, T. & Jentsch, S. A protein translocation defect linked to ubiquitin conjugation at the endoplasmic reticulum. Nature 365, 176–179, doi:10.1038/365176a0 (1993).

5 Jensen, T. J. et al. Multiple proteolytic systems, including the proteasome, contribute to CFTR processing. Cell 83, 129–135, doi:10.1016/0092-8674(95)90241-4 (1995).

6 Ward, C. L., Omura, S. & Kopito, R. R. Degradation of CFTR by the ubiquitin-proteasome pathway. Cell 83, 121–127 (1995).

7 McCracken, A. A. & Brodsky, J. L. Assembly of ER-associated protein degradation in vitro: dependence on cytosol, calnexin, and ATP. J Cell Biol 132, 291–298, doi:10.1083/jcb.132.3.291 (1996).

8 Hampton, R. Y., Gardner, R. G. & Rine, J. Role of 26S proteasome and HRD genes in the degradation of 3-hydroxy-3-methylglutaryl-CoA reductase, an integral endoplasmic reticulum membrane protein. Mol Biol Cell 7, 2029–2044, doi:10.1091/mbc.7.12.2029 (1996).

9 Bordallo, J., Plemper, R. K., Finger, A. & Wolf, D. H. Der3p/Hrd1p is required for endoplasmic reticulum-associated degradation of misfolded lumenal and integral membrane proteins. Mol Biol Cell 9, 209–222, doi:10.1091/mbc.9.1.209 (1998).

10 Biunno, I. et al. Isolation of a pancreas-specific gene located on human chromosome 14q31: expression analysis in human pancreatic ductal carcinomas. Genomics 46, 284–286, doi:10.1006/geno.1997.5018 (1997).

11 Kaneko, M., Ishiguro, M., Niinuma, Y., Uesugi, M. & Nomura, Y. Human HRD1 protects against ER stress-induced apoptosis through ER-associated degradation. FEBS Lett 532, 147–152, doi:10.1016/s0014-5793(02)03660-8 (2002).

12 Hwang, J. & Qi, L. Quality Control in the Endoplasmic Reticulum: Crosstalk between ERAD and UPR pathways. Trends Biochem Sci 43, 593–605, doi:10.1016/j.tibs.2018.06.005 (2018).

13 Wang, H. H. et al. Hypomorphic variants of SEL1L-HRD1 ER-associated degradation are associated with neurodevelopmental disorders. J Clin Invest 134, doi:10.1172/JCI170054 (2024).

14 Weis, D. et al. Biallelic Cys141Tyr variant of SEL1L is associated with neurodevelopmental disorders, agammaglobulinemia, and premature death. J Clin Invest 134, doi:10.1172/JCI170882 (2024).

15 Qi, L., Tsai, B. & Arvan, P. New Insights into the Physiological Role of Endoplasmic Reticulum-Associated Degradation. Trends Cell Biol 27, 430–440, doi:10.1016/j.tcb.2016.12.002 (2017).

16 Bhattacharya, A. & Qi, L. ER-associated degradation in health and disease - from substrate to organism. J Cell Sci 132, jcs232850, doi:10.1242/jcs.232850 (2019).

17 Bays, N. W., Gardner, R. G., Seelig, L. P., Joazeiro, C. A. & Hampton, R. Y. Hrd1p/Der3p is a membrane-anchored ubiquitin ligase required for ER-associated degradation. Nat Cell Biol 3, 24–29, doi:10.1038/35050524 (2001).

18 Gardner, R. G. et al. Endoplasmic reticulum degradation requires lumen to cytosol signaling. Transmembrane control of Hrd1p by Hrd3p. J Cell Biol 151, 69–82, doi:10.1083/jcb.151.1.69 (2000).

19 Sun, S. et al. Sel1L is indispensable for mammalian endoplasmic reticulum-associated degradation, endoplasmic reticulum homeostasis, and survival. Proc Natl Acad Sci U S A 111, E582–591, doi:10.1073/pnas.1318114111 (2014).

20 Mueller, B., Lilley, B. N. & Ploegh, H. L. SEL1L, the homologue of yeast Hrd3p, is involved in protein dislocation from the mammalian ER. J Cell Biol 175, 261–270, doi:10.1083/jcb.200605196 (2006).

21 Mueller, B., Klemm, E. J., Spooner, E., Claessen, J. H. & Ploegh, H. L. SEL1L nucleates a protein complex required for dislocation of misfolded glycoproteins. Proc Natl Acad Sci USA 105, 12325–12330, doi:10.1073/pnas.0805371105 (2008).

22 Bhamidipati, A., Denic, V., Quan, E. M. & Weissman, J. S. Exploration of the topological requirements of ERAD identifies Yos9p as a lectin sensor of misfolded glycoproteins in the ER lumen. Mol Cell 19, 741–751, doi:10.1016/j.molcel.2005.07.027 (2005).

23 Kim, W., Spear, E. D. & Ng, D. T. Yos9p detects and targets misfolded glycoproteins for ER-associated degradation. Mol Cell 19, 753–764, doi:10.1016/j.molcel.2005.08.010 (2005).

24 Szathmary, R., Bielmann, R., Nita-Lazar, M., Burda, P. & Jakob, C. A. Yos9 protein is essential for degradation of misfolded glycoproteins and may function as lectin in ERAD. Mol Cell 19, 765–775, doi:10.1016/j.molcel.2005.08.015 (2005).

25 Christianson, J. C., Shaler, T. A., Tyler, R. E. & Kopito, R. R. OS-9 and GRP94 deliver mutant alpha1-antitrypsin to the Hrd1-SEL1L ubiquitin ligase complex for ERAD. Nat Cell Biol 10, 272–282, doi:10.1038/ncb1689 (2008).

26 Hosokawa, N. et al. Human XTP3-B forms an endoplasmic reticulum quality control scaffold with the HRD1-SEL1L ubiquitin ligase complex and BiP. J Biol Chem 283, 20914–20924, doi:10.1074/jbc.M709336200 (2008).

27 Lilley, B. N. & Ploegh, H. L. Multiprotein complexes that link dislocation, ubiquitination, and extraction of misfolded proteins from the endoplasmic reticulum membrane. Proc Natl Acad Sci U S A 102, 14296–14301, doi:10.1073/pnas.0505014102 (2005).

28 Ye, Y., Shibata, Y., Yun, C., Ron, D. & Rapoport, T. A. A membrane protein complex mediates retro-translocation from the ER lumen into the cytosol. Nature 429, 841–847, doi:10.1038/nature02656 (2004).

29 Ye, Y. et al. Inaugural Article: Recruitment of the p97 ATPase and ubiquitin ligases to the site of retrotranslocation at the endoplasmic reticulum membrane. Proc Natl Acad Sci USA 102, 14132–14138, doi:10.1073/pnas.0505006102 (2005).

30 Carvalho, P., Goder, V. & Rapoport, T. A. Distinct ubiquitin-ligase complexes define convergent pathways for the degradation of ER proteins. Cell 126, 361–373, doi:10.1016/j.cell.2006.05.043 (2006).

31 Schoebel, S. et al. Cryo-EM structure of the protein-conducting ERAD channel Hrd1 in complex with Hrd3. Nature 548, 352–355, doi:10.1038/nature23314 (2017).

32 Wu, X. et al. Structural basis of ER-associated protein degradation mediated by the Hrd1 ubiquitin ligase complex. Science 368, doi:10.1126/science.aaz2449 (2020).

33 Hwang, J. et al. Characterization of protein complexes of the endoplasmic reticulum-associated degradation E3 ubiquitin ligase Hrd1. J Biol Chem 292, 9104–9116, doi:10.1074/jbc.M117.785055 (2017).

34 Huang, C. H., Hsiao, H. T., Chu, Y. R., Ye, Y. & Chen, X. Derlin2 protein facilitates HRD1-mediated retro-translocation of sonic hedgehog at the endoplasmic reticulum. J Biol Chem 288, 25330–25339, doi:10.1074/jbc.M113.455212 (2013).

35 Huang, C. H., Chu, Y. R., Ye, Y. & Chen, X. Role of HERP and a HERP-related protein in HRD1-dependent protein degradation at the endoplasmic reticulum. J Biol Chem 289, 4444–4454, doi:10.1074/jbc.M113.519561 (2014).

36 Lin, L. L. et al. SEL1L-HRD1 interaction is required to form a functional HRD1 ERAD complex. Nat Commun 15, 1440, doi:10.1038/s41467-024-45633-0 (2024).

37 van der Goot, A. T., Pearce, M. M. P., Leto, D. E., Shaler, T. A. & Kopito, R. R. Redundant and Antagonistic Roles of XTP3B and OS9 in Decoding Glycan and Non-glycan Degrons in ER-Associated Degradation. Mol Cell 70, 516–530 e516, doi:10.1016/j.molcel.2018.03.026 (2018).

38 Schulz, J. et al. Conserved cytoplasmic domains promote Hrd1 ubiquitin ligase complex formation for ER-associated degradation (ERAD). J Cell Sci 130, 3322–3335, doi:10.1242/jcs.206847 (2017).

39 Shi, G. et al. ER-associated degradation is required for vasopressin prohormone processing and systemic water homeostasis. J Clin Invest 127, 3897–3912 (2017).

40 Kikkert, M. et al. Human HRD1 is an E3 ubiquitin ligase involved in degradation of proteins from the endoplasmic reticulum. J Biol Chem 279, 3525–3534, doi:10.1074/jbc.M307453200 (2004).

41 Abramson, J. et al. Accurate structure prediction of biomolecular interactions with AlphaFold 3. Nature 630, 493–500, doi:10.1038/s41586-024-07487-w (2024).

42 Pisa, R. & Rapoport, T. A. Disulfide-crosslink analysis of the ubiquitin ligase Hrd1 complex during endoplasmic reticulum-associated protein degradation. J Biol Chem 298, 102373, doi:10.1016/j.jbc.2022.102373 (2022).

43 Hwang, C., Sinskey, A. J. & Lodish, H. F. Oxidized redox state of glutathione in the endoplasmic reticulum. Science 257, 1496–1502, doi:10.1126/science.1523409 (1992).

44 Pleiner, T. et al. Structural basis for membrane insertion by the human ER membrane protein complex. Science 369, 433–436, doi:10.1126/science.abb5008 (2020).

45 Bai, L., You, Q., Feng, X., Kovach, A. & Li, H. Structure of the ER membrane complex, a transmembrane-domain insertase. Nature 584, 475–478, doi:10.1038/s41586-020-2389-3 (2020).

46 Jomaa, A. et al. Structure of the quaternary complex between SRP, SR, and translocon bound to the translating ribosome. Nature communications 8, 15470, doi:10.1038/ncomms15470 (2017).

47 Jomaa, A. et al. Molecular mechanism of cargo recognition and handover by the mammalian signal recognition particle. Cell reports 36, 109350, doi:10.1016/j.celrep.2021.109350 (2021).

48 Rao, B. et al. The cryo-EM structure of the human ERAD retrotranslocation complex. Sci Adv 9, eadi5656, doi:10.1126/sciadv.adi5656 (2023).

