## Supplemental Information for "Structural basis and pathological implications of the dimeric OS9-SEL1L-HRD1 ERAD Core Complex"

**This PDF file includes:**

Materials and Methods

Figures. S1 to S9

Tables S1 to S4

References

**MATERIALS AND METHODS**

**Cell lines**

HEK293T cells (ATCC) were cultured at 37°C with 5% CO_2_ in DMEM (Gibco) with 10% fetal bovine serum (Fisher Scientific). HEK293F cells (Gibco) were cultured at 37°C with 5% CO_2_ in Expi293™ Expression Medium (Gibco) with shaking.

**Protein expression and purification**

0.6 mg plasmid was transfected to 0.5 L HEK293F cells by 1.6 mg PEI Max (Polysciences) for protein expression. The medium was collected at 72 h post-transfection for protein purification. The pellet from 500 mL cells was resuspended in 30 mL buffer A (20 mM HEPES, 150 mM NaCl, pH7.5) supplemented by protease inhibitor (Sigma) and phosphatase inhibitor cocktail (Sigma) and disrupted by sonication. After 3500 x g centrifugation for 5 min, the resulting supernatant was then incubated with 1% (w/v) lauryl maltose neopentyl glycol (LMNG, Anatrace) at 4 ^o^C for 2 h. The insoluble fraction was removed by centrifugation (21,000 x g, 4 ^o^C, 30 min), and the supernatant was incubated with 1 mL anti-FLAG M2 resin (Sigma) at 4 ^o^C for 4 h. The beads were then washed with 20 column volume (CV) buffer B (20 mM HEPES, 150 mM NaCl, 0.01% LMNG, pH7.5) by gravity flow. Proteins were eluted in 5 CV buffer B supplemented by 0.1 mg/mL 3xFLAG peptide (APExBIO). The eluate was concentrated and further purified by size-exclusion chromatography (SEC) (Superose 6 Increase, 10/300 GL column, Cytiva) in buffer C (20 mM HEPES, 150 mM NaCl, 0.02% glyco-diosgenin/GDN, pH7.5) for cryo-EM.

**Cryo-EM sample preparation and data acquisition**

Protein samples were concentrated using 100 kDa filter (Amino) to 4 mg/mL for grid preparation. Aliquots of 3 μl protein samples in detergent were applied to Quantifoil R1.2/1.3 400 mesh Cu holy carbon grids (Quantifoil) glow discharged with amylamine, blotted for 8s at 4 °C and 100%, then plunge-frozen in liquid ethane using a Vitrobot Mark IV (FEI). Data was collected on the ThermoFisher Titan Krios equipped with a K3 direct electron detector. The microscope was operated at 300 kV at a nominal magnification of 130,000X with an energy filter slit width of 10 eV. A defocus range of -0.8 µm to -1.8 µm at 0.2 µm step sizes was used for data collection. Movies were collected at a dose of 60 e/Å^2^ across 40 frames per movie. A total of 6,350 movies were collected and the calibrated pixel size was 0.652 Å.

**Cryo-EM data processing**

Cryo-EM data was processed using CryoSPARC v4.6.0 ^1^. Movies were corrected for beam-induced motion and subjected to patch CTF estimation. Blob picker was initially used to select for particles with minimum and maximum diameters of 120 Å and 250 Å, respectively. 2D classification of the picked particles yielded classes with clear membrane-embedded protein domain elements at an approximate diameter of ~130 Å. These classes were used to generate templates and improve particle picking. Template picker was used to select for particles with a 200 Å diameter using the generated templates from selected 2D class averages. From the motion-corrected micrographs, 1,101,567 particles were extracted at a box size of 488x488 pixels. An initial 2D classification was conducted to select a subset of well-aligned particles from 2D class averages showing clear protein domain and secondary structure elements resolved within and outside the micelle. An initial model for the ERAD complex was then generated using ab-initio reconstruction from 163,759 selected particles.

Subsequent 3D classifications were performed to recover additional ERAD particles for 3D homogenous refinements. Starting with the initial 1,101,567 extracted particles, iterative heterogenous refinements were conducted to sort for ERAD particles and remove junk or poorly aligning particles. A final set of 192,786 particles were reconstructed using non-uniform refinement applying C2 symmetry. The average resolution of the final map was 3.64 Å, as determined by the gold standard Fourier Shell Correlation (FSC) from two independently processed half-particle sets at an FSC threshold of 0.143. A mask was then created for the SEL1L and OS9 domains visible in the cryo-EM map. Symmetry expansion, followed by local 3D refinement focused on SEL1L-OS9 using the generated mask, was conducted to overcome symmetry-breaking features such as subunit flexibility between the individual components in the ERAD dimer and improve the map resolution in the OS9-SEL1L interaction interface to 3.30 Å. Local resolution of the focused and unfocused map was determined in cryoSPARC at an FSC threshold of 0.143.

**Model building and refinement**

The predicted Alphafold 3 ^2^ model of OS9, SEL1L, and HRD1 was docked as a rigid body into the cryo-EM map using ChimeraX. Regions that were not resolved in the EM maps were removed from fitted model. We then broke the model into 5 rigid body elements: 2OS9, 2SEL1L, and the HRD1 dimer that were then adjusted into the cryo-EM map with COOT. The transmembrane domains of HRD1 and secondary structural elements of SEL1L and OS9 were further rotated and positioned in the density map based on visible side chains in COOT. The final model was iteratively real space refined in PHENIX ^3^ (version 1.20.1) with 3 macrocycles applying Ramachandran and secondary structure restraints, and non-crystallographic symmetry constraints. Molprobity was used to validate the model and resolve clashes. All model and map figures were made with ChimeraX.

**CRISPR/Cas9-based knockout (KO) HEK293T cells**

To generate SEL1L-, HRD1-, and FAM8A1- deficient HEK293T cells, sgRNA oligonucleotides designed for human *SEL1L* (5’-GGCTGAACAGGGCTATGAAG-3’), human *HRD1* (5’-GGACAAAGGCCTGGATGTAC-3’) or *FAM8A1* (gRNA1: 5’-GCGCGGCGGCTCCAATTTGT-3’, gRNA2: 5’-TTCGGCCTGGGGGTCGTCGC-3’) was inserted into lentiCRISPR v2 (plasmid 52961; Addgene). HEK293T cells grown in 10 cm petri dishes were transfected with indicated plasmids using 5 μl 1 mg/ml polyethylenimine (PEI, Sigma) per 1μg of plasmids for HEK293T cells. Cells were cultured 24 hours after transfection in medium containing 2 µg/ml puromycin for 24 hours and then in normal growth media.

**Plasmids**

The following plasmids were used in the study:

*HRD1* cDNA with a C-terminal FLAG tag was cloned from HEK293T cDNA and inserted into the pcDNA3 to generate pcDNA3-hHRD1(WT)-FLAG. The HRD1 mutant C2A (C291A/C294A), T93C, V94C, F95C, R96C, D97C, T93A, T93F and A91D were generated using the plasmid pcDNA-hHRD1(WT)-FLAG as template. HRD1 T93A/R96C, T93F/R96C, and A91D/R96C mutants were generated using the plasmid pcDNA-hHRD1(R96C)-FLAG as template. *hOS9* cDNA was cloned from HEK293T cDNA and inserted into the pcDNA3 to generate pcDNA3-hOS9(WT). The OS9 mutant D135C was generated using plasmid pcDNA3-hOS9(WT) as the template. *SEL1L* cDNA, with/without a C-terminal FLAG tag was cloned into the pcDNA3 to generate pcDNA3-SEL1L(WT)-FLAG and pcDNA3-SEL1L(WT). The SEL1L-FLAG mutants S658P and G585D were generated using the plasmid pcDNA-SEL1L(WT)-FLAG as the template. The double mutations of SEL1L S658P/G585D was generated using the plasmid pcDNA-SEL1L(S658P)-FLAG as the template. The SEL1L mutants D439C, M440C and N442C were generated using the plasmid pcDNA-SEL1L(WT) as the template. The OS9 mutant D135C was generated using plasmid pcDNA3-hOS9(WT) as the template. h*-proAVP(G57S)* with 1xStrep/10xHis tag and KDEL in its C-terminal was cloned from the plasmid pcDNA3-h-proAVP(G7S)-HA, which was described previously ^4^, to generate pcDNA3-h-proAVP(G57S)-Strep-KDEL and proAVP(G57S)-His-KDEL. All plasmids were validated by DNA sequencing.

**Western blot and antibodies (reducing SDS-PAGE)**

HEK293T or HEK293F cells were harvested and snap-frozen in liquid nitrogen. The proteins were extracted by sonication in Western blot lysis buffer (50 mM Tris-HCl at pH7.5, 150 mM NaCl, 1% Triton X-100, 1 mM EDTA) with protease inhibitor (Sigma) and phosphatase inhibitor cocktail (Sigma). Lysates were incubated on ice for 15 min and centrifuged at 16,000 g for 10 min. Supernatants were collected and analyzed for protein concentration using the Bio-Rad Protein Assay Dye (Bio-Rad). 20-50 μg of protein were denatured at 95°C for 5 min or 37°C for 1 hr in 5x SDS sample buffer (250 mM Tris-HCl pH 6.8, 10% sodium dodecyl sulfate, 0.05% Bromophenol blue, 50% glycerol, and 1.44 M β-mercaptoethanol). Protein was separated on SDS-PAGE, followed by electrophoretic transfer to PVDF (Fisher Scientific) membrane. The blots were incubated in 2% BSA/Tri-buffered saline tween-20 (TBST) with primary antibodies overnight at 4°C: anti-HSP90 (Santa Cruz, #sc-13119, 1:5,000), anti-GAPDH (Proteintech, #60004-1, 1:5000), anti-SEL1L (home-made, 1:10,000) ^5^, anti-HRD1 (Proteintech, #13473-1, 1:2,000), anti-OS9 (Abcam, #ab109510, 1:5,000), anti-CD147 (Proteintech, #11989-1, 1:3,000), anti-IRE1α (Cell Signaling, #3294, 1:2,000), anti-UBE2J1 (Santa Cruz, #sc-377002, 1:3,000), anti-DERL2 (gift, 1:1,000), anti-FLAG (Sigma, #F1804, 1:1,000), anti-HERPUD1 (HERP1) (Abcam, #ab150424, 1:3000), anti-FAM8A1 (Proteintech, #24746-1-AP,1:3000), anti-ubiquitin (Santa Cruz, #sc-8017, 1:1000), anti-His (Genscript, # A00174, 1:2000), anti-Strep (Sigma, # SAB2702216, 1:1000). Membranes were washed with TBST and incubated with secondary antibodies, HRP conjugated (Bio-Rad, 1:10,000) at room temperature for 1h for ECL chemiluminescence detection system (Bio-Rad) development. Band intensity was determined using Image lab (Bio-Rad) software.

**Mass spectrometry**

SEL1L- and HRD1- IP-MS in HEK293T cells have been described previously ^6^. Briefly, IP of endogenous SEL1L or HRD1 in WT, *SEL1L^-/-^* or *HRD1^-/-^* HEK293T cells were performed using 10 mg proteins from each sample lysed with the IP buffer (150 mM NaCl, 0.2% NP-40, 0.1% Triton X-100, 25 mM Tris-HCl pH7.5) supplemented with protease inhibitors, protein phosphatase inhibitors, and 10 mM N-ethylmaleimide. The cell lysates were first incubated anti-SEL1L (home-made) or anti-HRD1 (home-made or Cell Signaling #14773) at 4°C overnight, followed by incubation with Protein A agarose (Invitrogen, #20333) at 4°C for 2 hours. An IP reaction with IgG using cell lysate of *SEL1L^-/-^* *or HRD1^-/-^* cells was included as a negative control for SEL1L- or HRD1-IP, respectively.

The mass spectrometry was conducted by the Proteomics Resource Facility at the University of Michigan Medical School. The beads were resuspended in 50 µl of 0.1 M ammonium bicarbonate buffer (pH~8). Cysteines were reduced by adding 50 µl of 10 mM DTT and incubating at 45° C for 30 min. Samples were cooled to room temperature and alkylation of cysteines was achieved by incubating with 65 mM 2-Chloroacetamide, under darkness, for 30 min at room temperature. An overnight digestion with 1 µg sequencing-grade modified trypsin was carried out at 37° C with constant shaking in a Thermomixer. Digestion was stopped by acidification and peptides were desalted using SepPak C18 cartridges using manufacturer’s protocol (Waters). Samples were completely dried using vacufuge. Resulting peptides were dissolved in 0.1% formic acid/2% acetonitrile solution and were resolved on a nano-capillary reverse phase column (Acclaim PepMap C18, 2 micron, 50 cm, ThermoScientific) using a 0.1% formic acid/2% acetonitrile (Buffer A) and 0.1% formic acid/95% acetonitrile (Buffer B) gradient at 300 nl/min over a period of 180 min (2-25% buffer B in 110 min, 25-40% in 20 min, 40-90% in 5 min followed by holding at 90% buffer B for 10 min and re-equilibration with Buffer A for 30 min). Eluent was directly introduced into Q exactive HF mass spectrometer (Thermo Scientific, San Jose CA) using an EasySpray source. MS1 scans were acquired at 60K resolution (AGC target=3x106; max IT=50 ms). Data-dependent collision induced dissociation MS/MS spectra were acquired using Top speed method (3 seconds) following each MS1 scan (NCE ~28%; 15K resolution; AGC target 1x105; max IT 45 ms). Proteins were identified by searching the MS/MS data against UniProt entries using Proteome Discoverer (v2.4, Thermo Scientific). Search parameters included MS1 mass tolerance of 10 ppm and fragment tolerance of 0.2 Da; two missed cleavages were allowed; carbamidomethylation of cysteine was considered fixed modification and oxidation of methionine, deamidation of asparagine and glutamine were considered as potential modifications. False discovery rate (FDR) was determined using Percolator and proteins/peptides with an FDR of ≤1% were retained for further analysis.

SEL1L- or HRD1- interacting proteins were selected based on the peptide spectrum matches (PSMs) from the label-free IP-MS results. For each protein hit, the PSM value from the IgG must be 0; the PSMs ratio of the bait KO sample (negative control) to WT sample must be smaller than the ratio of the corresponding bait. Proteomic data has been deposited into a public database PRIDE (PXD043674 and PXD041882).

**Native disulfide crosslinking (non-reducing SDS-PAGE)**

HEK293T or HEK293F cells transfected with the indicated OS9, SEL1L, or HRD1 variants were harvested and snap-frozen in liquid nitrogen. Cell pellets were resuspended in 1x non-reducing SDS loading buffer (50 mM Tris-HCl pH6.8, 2% sodium dodecyl sulfate, 0.01% Bromophenol blue, 10% glycerol), and proteins were extracted by sonication. Lysates were incubated at room temperature for 15 min, followed by centrifugation at 16,000 x g for 10 min. The resulting supernatants were collected for SDS-PAGE and immunoblot analysis.

**DSP crosslinking (non-reducing SDS-PAGE)**

HEK293T cells transfected with the indicated HRD1 variants were treated with 2mM DSP crosslinker (Thermo Science) or DMSO at room temperature for 2 h. The crosslinking reaction was quenched by adding 20 mM Tris-HCl (pH7.5). Treated cells were then harvested and snap-frozen in liquid nitrogen. Cell pellets were resuspended in 1x non-reducing SDS loading buffer (50 mM Tris-HCl pH6.8, 2% sodium dodecyl sulfate, 0.01% Bromophenol blue, 10% glycerol), and proteins were extracted by sonication. Lysates were incubated at room temperature for 15 min, followed by centrifugation at 16,000 x g for 10 min. The resulting supernatants were collected for SDS-PAGE and immunoblot analysis.

**Co-immunoprecipitation (Co-IP) in HEK293T cells**

HEK293T were snap-frozen in liquid nitrogen and whole cell lysate was prepared in IP buffer (150 mM NaCl, 0.2% NP-40, 0.1% Triton X-100, 25 mM Tris-HCl pH 7.5) for anti-SEL1L-FLAG or anti-HRD1-FLAG IP supplemented with protease inhibitors, and protein phosphatase inhibitors. A total of ~5 mg protein lysates were incubated with 10 μl anti-FLAG agarose (Sigma, #A2220) antibody overnight at 4°C with gentle rocking. The incubated agaroses were washed three times with IP buffer and eluted in 0.1mg/ml 3xFLAG peptides. The resulting elution was incubated with 5x SDS sample buffer (250 mM Tris-HCl pH 6.8, 10% sodium dodecyl sulfate, 0.05% Bromophenol blue, 50% glycerol, and 1.44 M β-mercaptoethanol) at room temperature for 15min, followed by SDS-PAGE and immunoblot analysis.

**Denaturing immunoprecipitation for ubiquitination assay**

HEK293T cells were transfected with proAVP(G57S)-Strep and the indicated HRD1 variants. HEK293F cells were transfected with proAVP(G57S)-Strep, and OS9-SEL1L-HRD1 WT/C2A proteins. The transfected cells were treated with DMSO or 10 μM MG132 for 2 h. Treated cells were snap-frozen in liquid nitrogen and whole cell lysates was prepared in the Triton X-100 lysis buffer (50 mM Tris-HCl at pH7.5, 150 mM NaCl, 1% Triton X-100, 1 mM EDTA) with 1% SDS and 5 mM DTT, and denatured at 95°C for 10 min and centrifuged at 16,000×g for 10 min. Subsequently, supernatants were diluted 1:10 with NP-40 lysis buffer and incubated with 10 μl anti-Strep agarose (IBA, #6-6350) overnight at 4°C with gentle rocking. The incubated agaroses were washed three times with the Triton X-100 lysis buffer and eluted in 5 mM Biotin. The resulting elution was incubated with 5x SDS sample buffer (250 mM Tris-HCl pH 6.8, 10% sodium dodecyl sulfate, 0.05% Bromophenol blue, 50% glycerol, and 1.44 M β-mercaptoethanol) at 95^o^C for 5 min followed by SDS-PAGE and Immunoblot. The samples were loaded into a 6-15% gradient SDS-PAGE gel for separation.

**SEL1L and HRD1 disease variants**

SEL1L G585D and S658P have been reported previously ^6-8^. We recently identified through Lifera Omics Database a novel, disease-associated variant of HRD1, *HRD1 A91D*, in a one-year-old boy with congenital heart defects and early-onset pulmonary dysfunction, born to consanguineous parents.

**Statistics**

The cryo-EM data collection, refinement and validation statistics are shown in Table S4. All experiments have been repeated at least two to three times and/or performed with multiple independent biological samples from which representative data are shown.

**References**

1	Punjani, A., Rubinstein, J. L., Fleet, D. J. & Brubaker, M. A. cryoSPARC: algorithms for rapid unsupervised cryo-EM structure determination. *Nat Methods* **14**, 290-296, doi:10.1038/nmeth.4169 (2017).

2	Abramson, J. *et al.* Accurate structure prediction of biomolecular interactions with AlphaFold 3. *Nature* **630**, 493-500, doi:10.1038/s41586-024-07487-w (2024).

3	Liebschner, D. *et al.* Macromolecular structure determination using X-rays, neutrons and electrons: recent developments in Phenix. *Acta Crystallogr D Struct Biol* **75**, 861-877, doi:10.1107/S2059798319011471 (2019).

4	Shi, G. *et al.* ER-associated degradation is required for vasopressin prohormone processing and systemic water homeostasis. *J Clin Invest* **127**, 3897-3912, doi:10.1172/JCI94771 (2017).

5	Zhou, Z. *et al.* Endoplasmic reticulum-associated degradation regulates mitochondrial dynamics in brown adipocytes. *Science* **368**, 54-60, doi:10.1126/science.aay2494 (2020).

6	Lin, L. L. *et al.* SEL1L-HRD1 interaction is required to form a functional HRD1 ERAD complex. *Nat Commun* **15**, 1440, doi:10.1038/s41467-024-45633-0 (2024).

7	Kyostila, K. *et al.* A SEL1L mutation links a canine progressive early-onset cerebellar ataxia to the endoplasmic reticulum-associated protein degradation (ERAD) machinery. *PLoS Genet* **8**, e1002759, doi:10.1371/journal.pgen.1002759 (2012).

8	Wang, H. H. *et al.* Hypomorphic variants of SEL1L-HRD1 ER-associated degradation are associated with neurodevelopmental disorders. *J Clin Invest* **134**, doi:10.1172/JCI170054 (2024).

**SUPPLEMENTAL FIGURES**

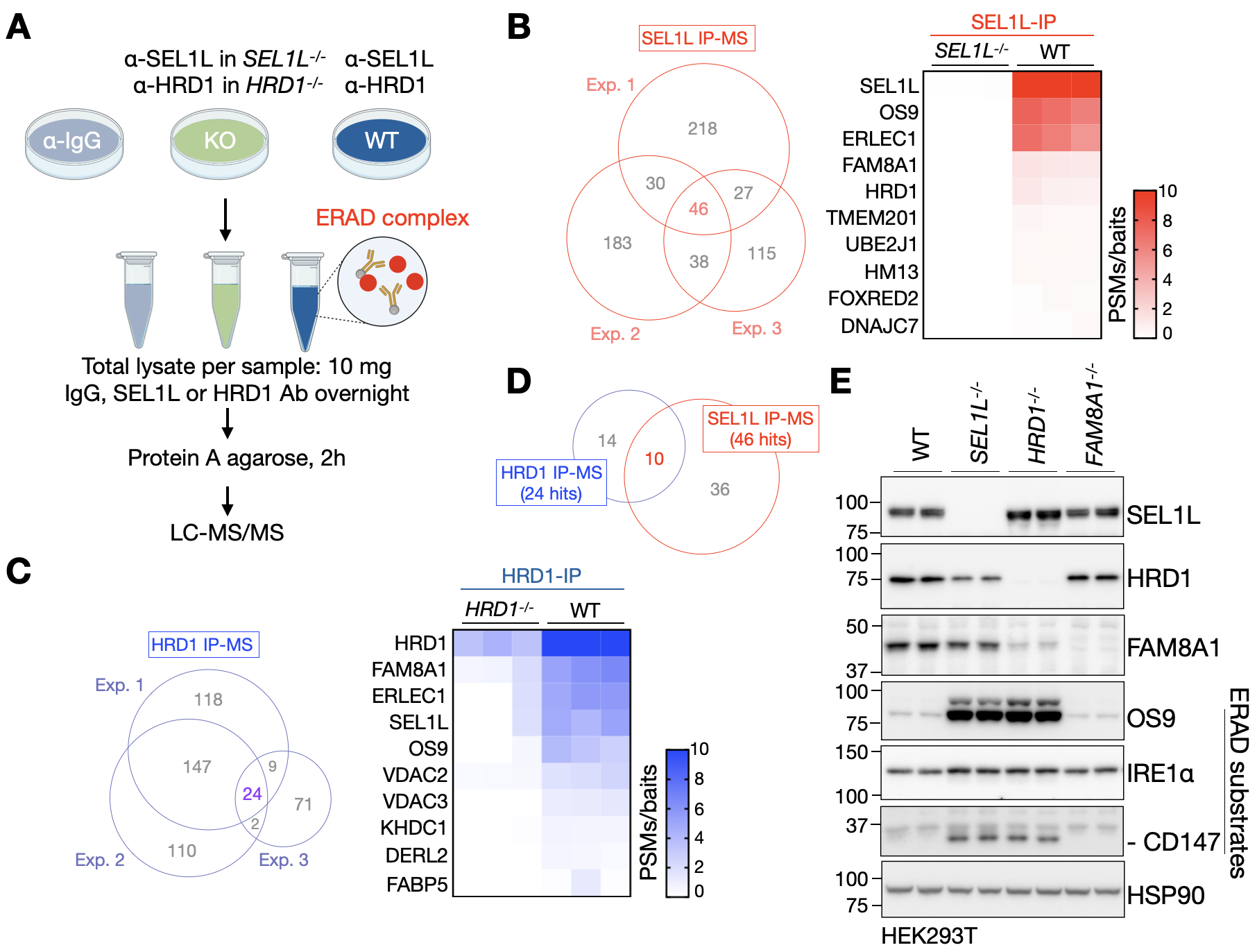

**Fig. S1 Identification of the core components of SEL1L-HRD1 ERAD machinery by IP-MS**

(**A**) Flow chart of IP-MS to detect endogenous SEL1L- or HRD1- interacting proteins in HEK293T.

(**B**) Venn diagram showing SEL1L-interacting proteins identified in three independent experiments (left) and the heatmap showing the top 10 hits of SEL1L-interacting proteins (right). Plotted values are peptide-spectrum matches (PSMs) normalized to their baits from three independent experiments. The result of 46 hits is shown in the Table S1.

(**C**) Venn diagram showing HRD1-interacting proteins identified in three independent experiments (left) and the heatmap showing the top 10 hits of HRD1-interacting proteins (right). Plotted values are peptide-spectrum matches (PSMs) normalized to their baits from three independent experiments. The result of 24 hits is shown in the Table S2.

(**D**) Venn diagram showing the overlapping hits of SEL1L- and HRD1- interacting proteins. Heatmap showing the top 5 hits is shown in Fig. 1A. The result of 10 overlapping hits is shown in Table S3.

(**E**) Western blot analysis of SEL1L, HRD1, FAM8A1 and known endogenous substrates (OS9, IRE1α, CD147) in WT, *SEL1L^-/-^*, *HRD1^-/-^* and *FAM8A1^-/-^* HEK293T cells (two independent repeats), showing dispensable role of FAM8A1 in HRD1 ERAD.

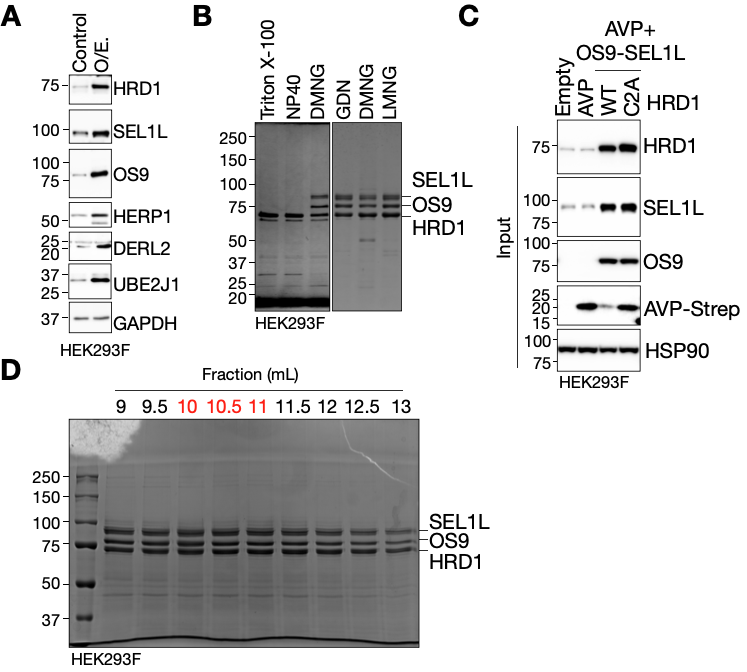

**Fig. S2 Purification and functional analysis of the OS9-SEL1L-HRD1 core complex**

(**A**) Western blot analysis of ERAD components in HEK293F cells expressing HRD1-FLAG, SEL1L, OS9, HERP1, DERL2 and UBE2J1, using GAPDH as loading control. O/E., overexpression.

(**B**) SDS-PAGE analysis (Coomassie blue staining) following HRD1-FLAG IP in HEK293F cells expressing ERAD components from (**A**) with different detergents for solubilization.

(**C**) Input of denaturing IP (Fig. 1B) of Strep-agarose in HEK293F cells expressing the ERAD substrate proAVP(G57S)-Strep, OS9, SEL1L, and HRD1-WT-FLAG or ligase-dead HRD1 mutant C2A (C291A/C293A)-FLAG, with 10 μM MG132 for 2 hours, to measure proAVP(G57S) ubiquitination (two independent repeats).

(**D**) SDS-PAGE analysis (Coomassie blue staining) of the fractions from Fig. 1C.

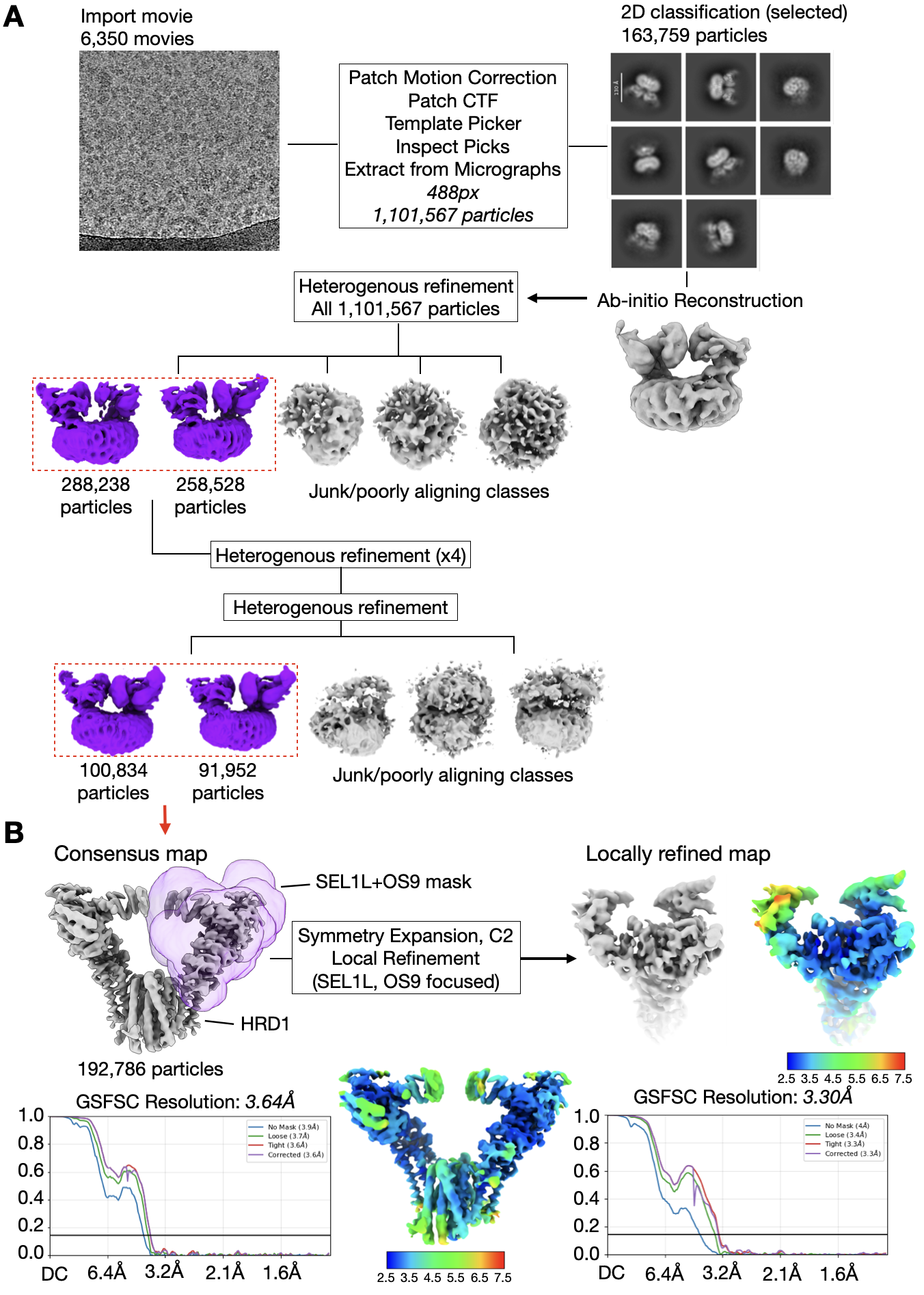

**Fig.S3 Single particle cryo-EM processing workflow used to determine structure of OS9-SEL1L-HRD1 ERAD complex**

(**A**) Collected movies were motion and CTF corrected in cryoSPARC prior to particle picking. Particles were extracted for 2D classification, and the selected particles were reconstructed to generate an initial structure of the ERAD complex. Heterogenous refinements on the full set of extracted particles was then used to iteratively remove junk or poorly aligning particles (grey) from ERAD particles (purple, red box). For clarity, representative outputs from the initial and final heterogeneous refinements are shown to illustrate the classification strategy.

(**B**) The final set of 192,786 particles were refined to an average resolution of 3.64 Å using gold standard Fourier Shell Correlation (GSFSC) at an FSC cutoff of 0.143. A mask was generated for the regions of SEL1L and OS9 resolved in our structure (purple transparent surface). Symmetry expansion of the particles followed by local refinement was conducted to improve the map quality in the OS9-SEL1L interaction interface. The local resolution was determined in cryoSPARC at FSC cutoff of 0.143. Unsharpened maps are shown for visualization.

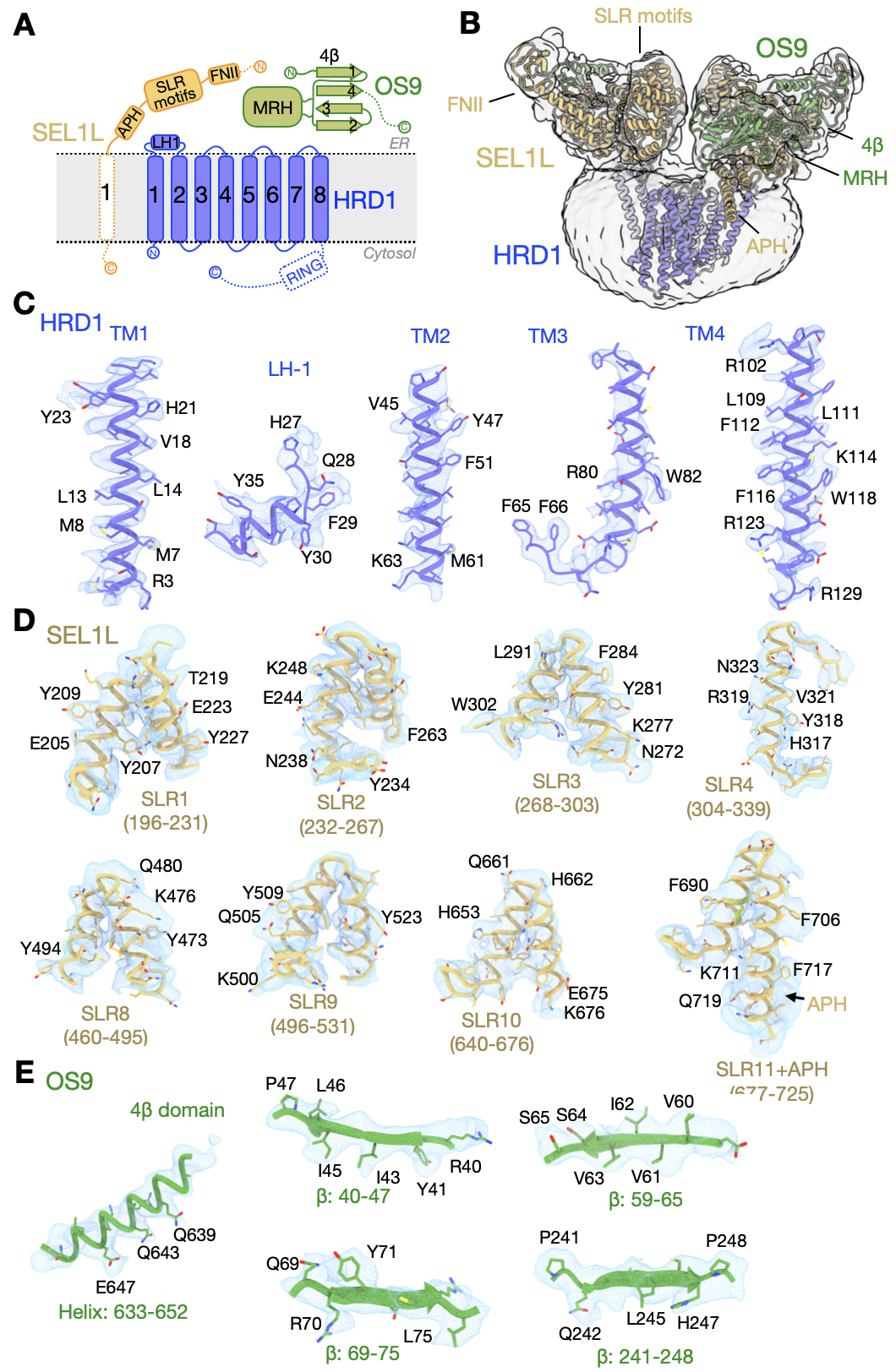

**Fig. S4 Schematic diagram and cryo-EM maps for the structural elements of OS9-SEL1L-HRD1 ERAD complex**

(**A**) Schematic diagram of the OS9-SEL1L-HRD1 ERAD complex showing their domains and motifs. The dotted lines and boxes indicate regions that were not resolved. APH, amphipathic helix; SLR, Sel-like repeat motifs, FNII, Fibronectin type II domain; RING, Really Interesting New Gene domain; LH1: loop helix between TM1-2 of HRD1; MRH, mannose 6-phosphate receptor homology domain; 4β, four β-sheet domain.

(**B**) Model of OS9-SEL1L-HRD1 fitted into cryo-EM map filtered to 6Å. Domains and motifs are labeled.

(**C-E**) Fits of the indicated model regions into the cryo-EM map. HRD1 is fitted into the consensus cryo-EM map filtered to 4Å (**C**). SEL1L and OS9 is fitted into locally refined cryo-EM map filtered to 4Å (**D-E**).

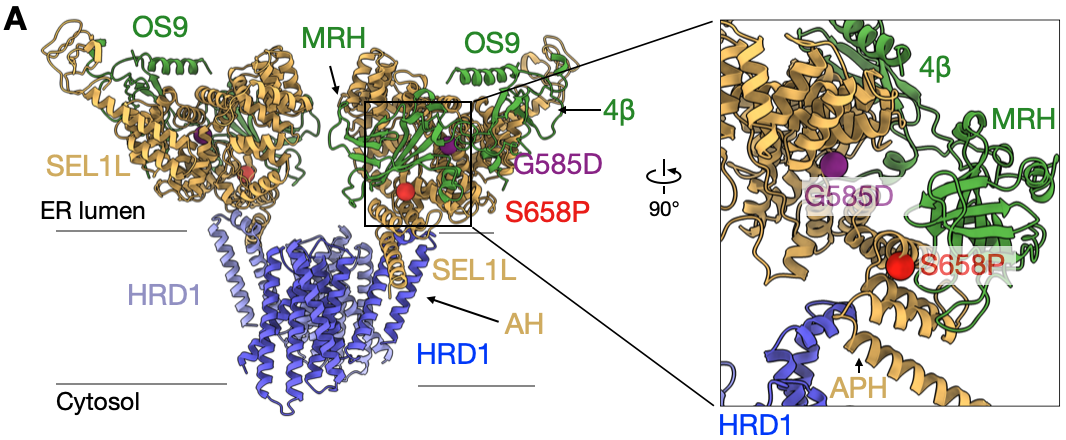

**Fig. S5 Location of two disease variants in the ERAD structure model**

(**A**) Side view of the models for the OS9-SEL1L-HRD1 complex, with two SEL1L disease variants highlighted in purple (G585D) and red (S658P). APH, amphipathic helix of SEL1L. MRH, mannose 6-phosphate receptor homology domain; 4β, four β-sheet domain.

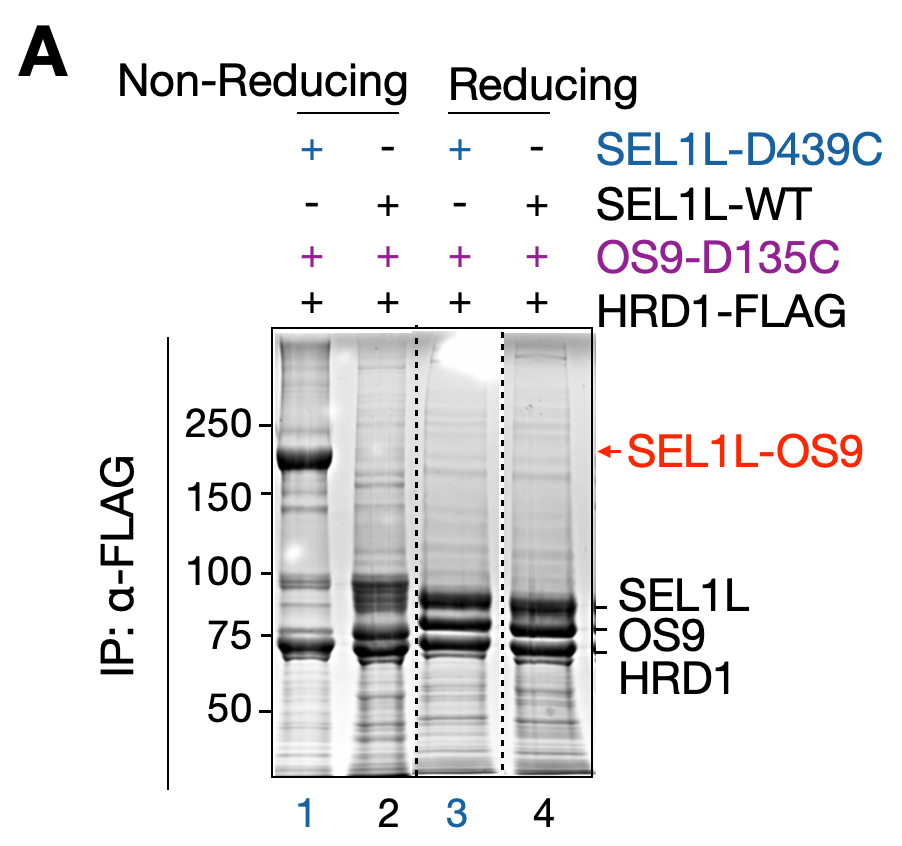

**Fig. S6 OS9-SEL1L dimerization *in vivo***

(**A**) Reducing and nonreducing SDS-PAGE analysis (Coomassie blue staining) of HRD1-FLAG IP in HEK293F cells expressing indicated OS9 and SEL1L mutations.

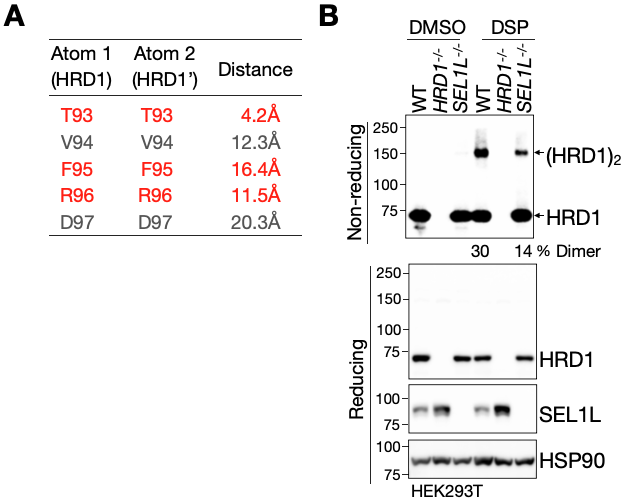

**Fig. S7 HRD1 dimerization *in vivo***

(**A**) Distance between the indicated HRD1 residues shown in Fig. 4B.

(**B**) Reducing and nonreducing SDS-PAGE and Western blot analysis of DSP-mediated crosslinking of HRD1 in WT and *HRD1^-/-^* HEK293T cells, following 2-hr 2mM DSP treatment. The percentage of the dimeric HRD1 in total HRD1 shown below the gel (two independent repeats).

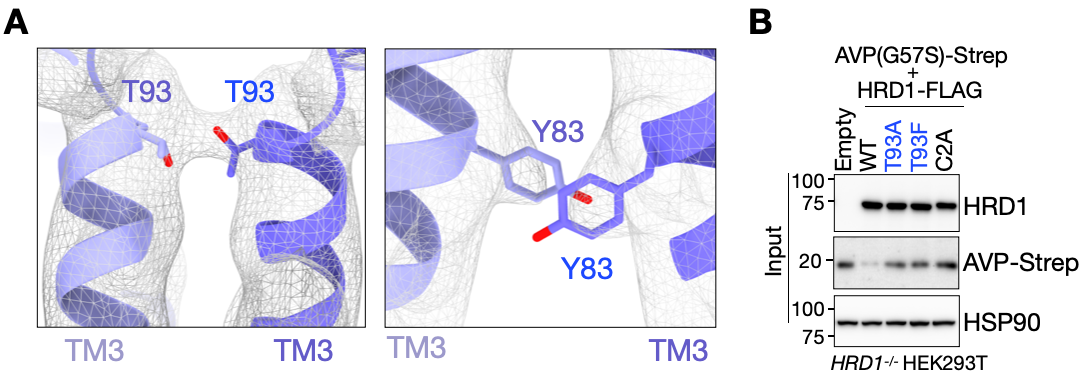

**Fig. S8 Interface of HRD1 dimer formed by TM3**

(**A**) Interactions of T93-T93, and Y83-Y83 in the HRD1 TM3 dimer interface (Fig. 5A). The cryo-EM map is filtered to 4Å and shown as a mesh.

(**B**) The input of denaturing IP experiment shown in Fig. 5E.

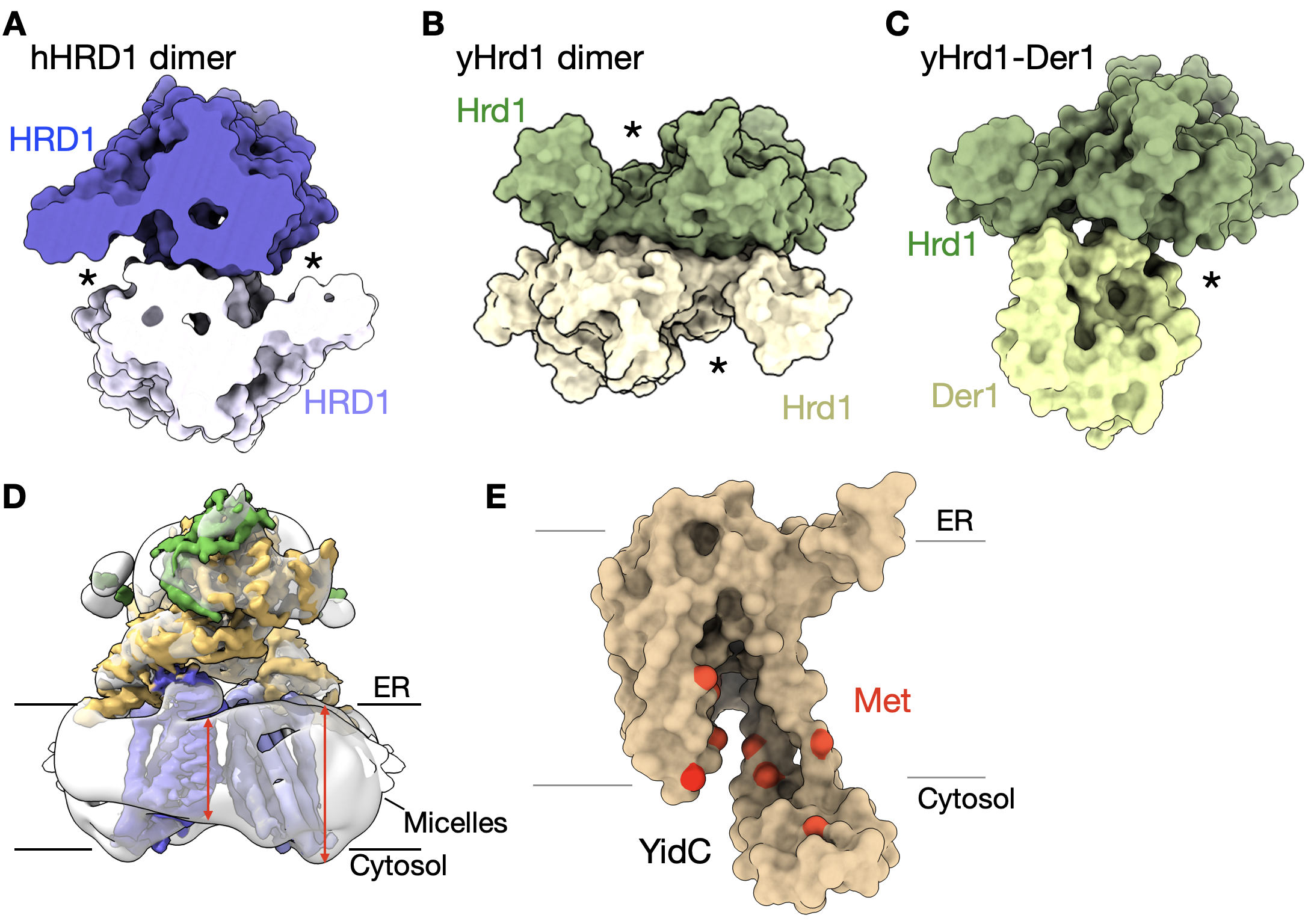

**Fig. S9 Putative substrate-conducting channels and membrane bending formed by HRD1 dimer, and methionine-rich bristles within YidC.**

(**A-C**) Surface model of the membrane-embedded region of human HRD1 (hHRD1) dimer (**A**), yeast Hrd1 (yeast) dimer (**B**), and yeast monomeric Hrd1-Der1 (**C**), viewed from the ER lumen. Asterisks indicate the putative substrate retrotranslocation channels.

(**D**) Cryo-EM map of the OS9-SEL1L-HRD1 complex, with the surrounding micelle in transparent gray. Distances in the micelle are shown in red.

(**E**) Surface model of the membrane-embedded region of the bacteria YidC (PDB: 3WO6). Methionine (Met) residues in the putative substrate retrotranslocation channel are highlighted in red, showing methionine-conducted bristles.

**Table S1. SEL1L-IP mass spectrometry result**

| No. | Protein | IgG | | | SEL1L-/- | | | WT | | |
| --- | --- | --- | --- | --- | --- | --- | --- | --- | --- | --- |
|  |  | 1st | 2nd | 3rd | 1st | 2nd | 3rd | 1st | 2nd | 3rd |
| 1 | SE1L1 | 0 | 0 | 0 | 4 | 1 | 2 | 260 | 341 | 304 |
| 2 | OS9 | 0 | 0 | 0 | 0 | 0 | 0 | 199 | 241 | 190 |
| 3 | ERLEC1 | 0 | 0 | 0 | 0 | 0 | 0 | 184 | 217 | 158 |
| 4 | FAM8A1 | 0 | 0 | 0 | 0 | 0 | 0 | 37 | 41 | 34 |
| 5 | HRD1 | 0 | 0 | 0 | 0 | 0 | 0 | 31 | 27 | 23 |
| 6 | TMEM201 | 0 | 0 | 0 | 0 | 0 | 0 | 13 | 13 | 12 |
| 7 | UBE2J1 | 0 | 0 | 0 | 0 | 0 | 0 | 10 | 14 | 11 |
| 8 | HM13 | 0 | 0 | 0 | 0 | 0 | 0 | 11 | 11 | 10 |
| 9 | FOXRED2 | 0 | 0 | 0 | 0 | 0 | 0 | 2 | 12 | 8 |
| 10 | DNAJC7 | 0 | 0 | 0 | 0 | 0 | 0 | 4 | 3 | 11 |
| 11 | HYOU1 | 0 | 0 | 0 | 0 | 0 | 0 | 5 | 5 | 4 |
| 12 | LRIG2 | 0 | 0 | 0 | 0 | 0 | 0 | 3 | 4 | 7 |
| 13 | HPSE | 0 | 0 | 0 | 0 | 0 | 0 | 3 | 8 | 3 |
| 14 | SIM14 | 0 | 0 | 0 | 0 | 0 | 0 | 3 | 6 | 3 |
| 15 | VDAC3 | 0 | 0 | 0 | 0 | 0 | 0 | 1 | 5 | 5 |
| 16 | PLOD2 | 0 | 0 | 0 | 0 | 0 | 0 | 6 | 2 | 2 |
| 17 | EDEM3 | 0 | 0 | 0 | 0 | 0 | 0 | 4 | 4 | 2 |
| 18 | EMILIN3 | 0 | 0 | 0 | 0 | 0 | 0 | 4 | 2 | 3 |
| 19 | TMEM9 | 0 | 0 | 0 | 0 | 0 | 0 | 4 | 2 | 3 |
| 20 | PRADC1 | 0 | 0 | 0 | 0 | 0 | 0 | 1 | 5 | 3 |
| 21 | CHST12 | 0 | 0 | 0 | 0 | 0 | 0 | 3 | 1 | 4 |
| 22 | PIGK | 0 | 0 | 0 | 0 | 0 | 0 | 2 | 3 | 3 |
| 23 | JAM3 | 0 | 0 | 0 | 0 | 0 | 0 | 2 | 3 | 3 |
| 24 | NCLN | 0 | 0 | 0 | 0 | 0 | 0 | 1 | 3 | 3 |
| 25 | NUP93 | 0 | 0 | 0 | 0 | 0 | 0 | 3 | 1 | 2 |
| 26 | YME1L1 | 0 | 0 | 0 | 0 | 0 | 0 | 3 | 1 | 2 |
| 27 | SUN2 | 0 | 0 | 0 | 0 | 0 | 0 | 3 | 2 | 1 |
| 28 | STT3A | 0 | 0 | 0 | 0 | 0 | 0 | 2 | 3 | 1 |
| 29 | LAMC1 | 0 | 0 | 0 | 0 | 0 | 0 | 2 | 2 | 2 |
| 30 | PIGS | 0 | 0 | 0 | 0 | 0 | 0 | 1 | 3 | 2 |
| 31 | TMED9 | 0 | 0 | 0 | 0 | 0 | 0 | 1 | 3 | 2 |
| 32 | TMEM30A | 0 | 0 | 0 | 0 | 0 | 0 | 3 | 1 | 1 |
| 33 | METTL2B | 0 | 0 | 0 | 0 | 0 | 0 | 3 | 1 | 1 |
| 34 | TMTC3 | 0 | 0 | 0 | 0 | 0 | 0 | 2 | 2 | 1 |
| 35 | TMEM33 | 0 | 0 | 0 | 0 | 0 | 0 | 1 | 2 | 2 |
| 36 | DDOST | 0 | 0 | 0 | 0 | 0 | 0 | 2 | 1 | 1 |
| 37 | LOX | 0 | 0 | 0 | 0 | 0 | 0 | 2 | 1 | 1 |
| 38 | CERS2 | 0 | 0 | 0 | 0 | 0 | 0 | 2 | 1 | 1 |
| 39 | PTK7 | 0 | 0 | 0 | 0 | 0 | 0 | 1 | 1 | 2 |
| 40 | CAND1 | 0 | 0 | 0 | 0 | 0 | 0 | 1 | 2 | 1 |
| 41 | GARS1 | 0 | 0 | 0 | 0 | 0 | 0 | 1 | 1 | 1 |
| 42 | CF120 | 0 | 0 | 0 | 0 | 0 | 0 | 1 | 1 | 1 |
| 43 | NUP133 | 0 | 0 | 0 | 0 | 0 | 0 | 1 | 1 | 1 |
| 44 | HERPUD1 | 0 | 0 | 0 | 0 | 0 | 0 | 1 | 1 | 1 |
| 45 | UBE2D3 | 0 | 0 | 0 | 0 | 0 | 0 | 1 | 1 | 1 |
| 46 | BAP1 | 0 | 0 | 0 | 0 | 0 | 0 | 1 | 1 | 1 |

46 hits for SEL1L interacting proteins from three independent repeats. The values in the table represent the total number of the sequenced peptides (PSMs value). The top 10 hits are highlighted in red.

**Table S2. HRD1-IP mass spectrometry result**

| No. | Protein | IgG | | | HRD1-/- | | | WT | | |
| --- | --- | --- | --- | --- | --- | --- | --- | --- | --- | --- |
|  |  | 1st | 2nd | 3rd | 1st | 2nd | 3rd | 1st | 2nd | 3rd |
| 1 | HRD1 | 0 | 0 | 0 | 47 | 47 | 33 | 135 | 109 | 94 |
| 2 | FAM8A1 | 0 | 0 | 0 | 8 | 8 | 19 | 70 | 66 | 62 |
| 3 | ERLEC1 | 0 | 0 | 0 | 0 | 0 | 18 | 66 | 62 | 54 |
| 4 | SEL1L | 0 | 0 | 0 | 0 | 0 | 17 | 65 | 45 | 49 |
| 5 | OS9 | 0 | 0 | 0 | 0 | 0 | 5 | 53 | 36 | 25 |
| 6 | VDAC2 | 0 | 0 | 0 | 4 | 4 | 4 | 23 | 21 | 23 |
| 7 | VDAC3 | 0 | 0 | 0 | 0 | 0 | 0 | 15 | 13 | 13 |
| 8 | KHDC1 | 0 | 0 | 0 | 0 | 0 | 1 | 9 | 7 | 6 |
| 9 | DERL2 | 0 | 0 | 0 | 0 | 0 | 0 | 8 | 6 | 3 |
| 10 | FABP5 | 0 | 0 | 0 | 0 | 0 | 0 | 1 | 14 | 1 |
| 11 | VDAC1 | 0 | 0 | 0 | 0 | 0 | 0 | 5 | 7 | 4 |
| 12 | UBE2D3 | 0 | 0 | 0 | 0 | 0 | 0 | 4 | 7 | 4 |
| 13 | SURF4 | 0 | 0 | 0 | 0 | 0 | 1 | 5 | 6 | 3 |
| 14 | HACD3 | 0 | 0 | 0 | 0 | 0 | 0 | 2 | 2 | 7 |
| 15 | OMA1 | 0 | 0 | 0 | 0 | 0 | 0 | 3 | 1 | 2 |
| 16 | STT3A | 0 | 0 | 0 | 0 | 0 | 0 | 3 | 3 | 1 |
| 17 | HM13 | 0 | 0 | 0 | 0 | 0 | 0 | 2 | 3 | 2 |
| 18 | UNC50 | 0 | 0 | 0 | 0 | 0 | 0 | 2 | 2 | 2 |
| 19 | UBE2D1 | 0 | 0 | 0 | 0 | 0 | 0 | 2 | 2 | 2 |
| 20 | NOP58 | 0 | 0 | 0 | 0 | 0 | 0 | 2 | 2 | 1 |
| 21 | CHERP | 0 | 0 | 0 | 0 | 0 | 0 | 2 | 2 | 1 |
| 22 | PRADC1 | 0 | 0 | 0 | 0 | 0 | 0 | 1 | 2 | 2 |
| 23 | TMUB1 | 0 | 0 | 0 | 0 | 0 | 0 | 1 | 2 | 1 |
| 24 | OSTC | 0 | 0 | 0 | 0 | 0 | 0 | 1 | 1 | 1 |

24 hits for HRD1 interacting proteins from three independent repeats. The values in the table represent the total number of the sequenced peptides (PSMs value). The top 10 hits are highlighted in blue.

**Table S3. SEL1L- and HRD1- interacting proteins**

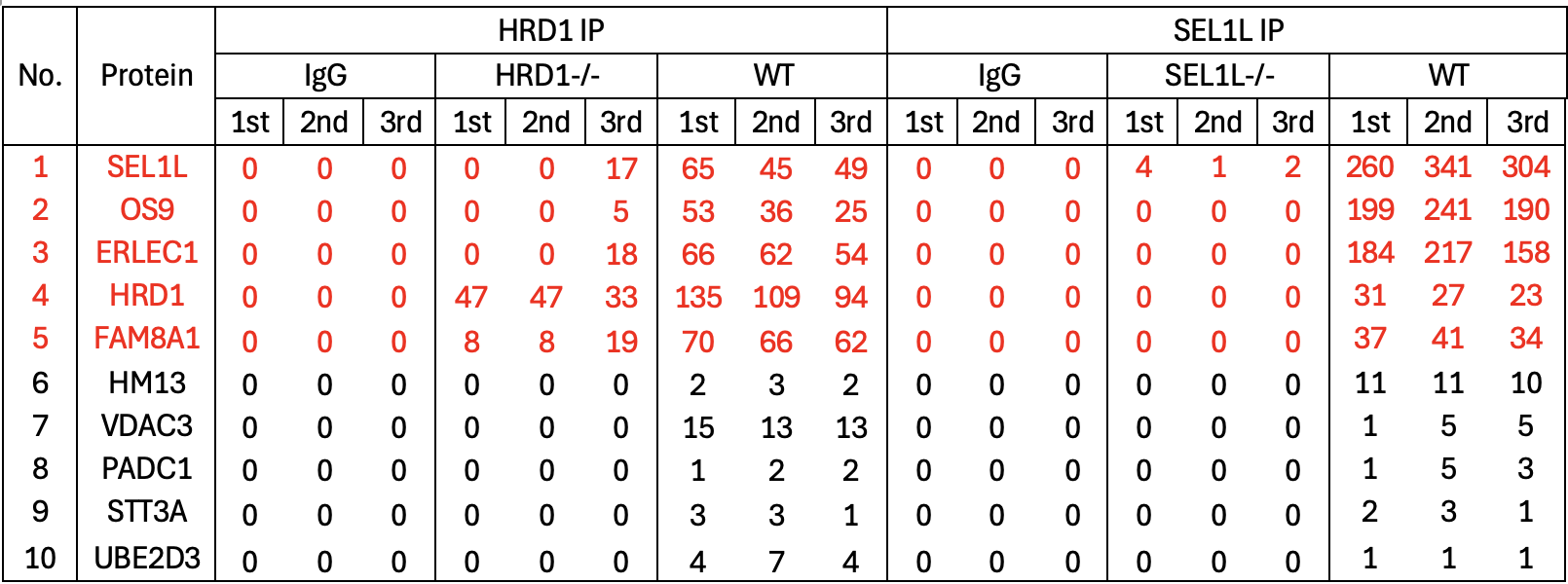

10 overlapping hits for SEL1L- and HRD1- interacting proteins from SEL1L- and HRD1- IP-MS results. The values in the table represent the total number of the sequenced peptides (PSMs value). The top 5 hits are highlighted in red.

**Table S4. Validation statistics for the cryo-EM data collection and model refinement.**

|  | | **OS9-SEL1L-HRD1** | **OS9-SEL1L (focused map)** |
| --- | --- | --- | --- |
| EMDB code | | EMD-70448 | EMD-70452 |
| PDB code | | 9OG0 |  |
| **Data collection and processing** | | |  |
| Nominal magnification | | 130,000x | 130,000x |
| Voltage (kV) | | 300 | 300 |
| Electron exposure (e^–^/Å^2^) | | 60 | 60 |
| Defocus range (μm) | | -1.8/-0.8 | -1.8/-0.8 |
| Pixel size (Å) | | 0.652 | 0.652 |
| Initial particle images (no.) | | 1,101,567 | 1,101,567 |
| Final particle images (no.) | | 192,786 | 192,786 |
| Map resolution at FSC=0.143 (Å) | | 3.64 | 3.30 |
| **Structure refinement in PHENIX 1.20.1** | | |  |
| Model resolution at FSC=0.5 (Å) | | 4.0 |  |
| CC_mask_ | | 0.61 |  |
| Map sharpening B factor (Å^2^) | | 101.2 |  |
| Model composition | | |  |
| Non-hydrogen atoms | | 17,976 |  |
| Protein residues | | 2248 |  |
| B factors min/max/mean (Å^2^) | | |  |
| Protein | | 1.90/178.93/80.02 |  |
| RMSD | | |  |
| Bond lengths (Å) | | 0.002 |  |
| Bond angles (°) | | 0.468 |  |
| **Validation** | | |  |
| MolProbity score | | 2.17 |  |
| Clashscore | | 7.81 |  |
| Rotamer Outliers (%) | | 3.16 |  |
| Ramachandran plot | | |  |
| Favored (%) | | 94.80 |  |
| Allowed (%) | | 5.06 |  |
| Outliers (%) | | 0.13 |  |
